# Lenacapavir binding to immature Gag triggers the emergence of giant HIV-1 virions

**DOI:** 10.1101/2025.07.16.665102

**Authors:** Wright Andrews Ofotsu Amesimeku, Yoshihiro Nakata, Nami Monde, Hiromi Terasawa, Hirotaka Ode, Hiroyuki Sasaki, Takeshi Matsui, Perpetual Nyame, Md. Jakir Hossain, Akatsuki Saito, Tomohiro Sawa, Terumasa Ikeda, Yosuke Maeda, Yasumasa Iwatani, Kazuaki Monde

**Affiliations:** Department of Microbiology, Faculty of Life Sciences, Kumamoto University, Kumamoto 860-8556, Japan; Department of Infectious Diseases and Immunology, Clinical Research Center, National Hospital Organization Nagoya Medical Center, 4-1-1 Sannomaru, Naka-ku, Nagoya 460-0001, Japan; Laboratory for Evolutionary Cell Biology of the Skin, School of Bioscience and Biotechnology, Tokyo University of Technology, 1404-1, Katakura-cho, Tokyo 192-0982, Japan; Department of Veterinary Medicine, University of Miyazaki, 1-1 Gakuen Kibanadai-nishi, Miyazaki 889-2192, Japan; Division of Molecular Virology and Genetics, Joint Research Center for Human Retrovirus Infection, Kumamoto University, Kumamoto 860-0811, Japan; Department of Nursing, Kibi International University, Takahashi, Okayama 716-8508, Japan; Collaboration Unit for Infection, Joint Research Center for Human Retrovirus Infection, Kumamoto University, Kumamoto, 860-0811, Japan

## Abstract

Lenacapavir (LEN), a potent capsid inhibitor, suppresses reverse transcription and nuclear import by disrupting capsid core formation in human immunodeficiency virus type 1 (HIV-1). However, its effect on the late-stage viral assembly remains unclear. Although p24 ELISA suggested that LEN inhibited HIV-1, both RT-dPCR and Vpr-HiBiT assays indicated no reduction in the viral output. This discrepancy was attributed to LEN-induced epitope masking of p24, which was reversed by altered sample processing. Furthermore, LEN induced numerous abnormal, discrete clusters of Gag at the plasma membrane. Mass photometry and transmission electron microscopy revealed heterogeneous viral-like particles (LENiVLPs; diameter >200 nm), containing both Gag and Env proteins. Although LENiVLPs retained membrane fusion capability, they were non-infectious owing to post-entry defects. Thus, LEN binds to the precursor Gag and promotes aberrant particle formation, offering novel insights that could guide the development of next-generation LEN-based therapeutics targeting the late stages of the viral life cycle.

## Main

In 2022, lenacapavir (LEN; SUNLENCA; formerly GS-6207; Gilead Sciences, Inc.) gained clinical approval as the first-in-class, small-molecule, long-acting human immunodeficiency virus type 1 (HIV-1) capsid (CA) inhibitor^1,2^. PF-3450074 (PF74) —with a binding pocket similar to LEN^3^—was developed previously^4^. However, it possesses weaker antiviral activity and greater cytotoxicity^4^. Similarly, GS-CA1—another compound targeting the same site— does exhibit picomolar potency, but it falls short of the enhanced potency and stability demonstrated by LEN^5^. Bevirimat (BVM, also known as PA-457), a maturation inhibitor acting on the six-helix bundle at the CA C-terminal domain (CTD) and spacer peptide 1 (SP1) of the HIV-1 Gag polyprotein^6,7^. Despite its potential, it was not approved because of the presence of resistance-associated mutations in patients carrying specific Gag polymorphisms^8^. These limitations have positioned LEN as the first approved anti-retroviral drug (ARV) that directly targets the HIV-1 Gag, particularly its CA domain^1,2^. LEN outperforms agents such as cabotegravir and rilpivirine in several respects: it requires only biannual dosing^9^, exhibits picomolar-level antiviral activity, lacks cross-resistance with existing ARVs, and operates through multiple inhibitory mechanisms^1^.

The HIV-1 CA protein critically influences the entire viral life cycle, making it a promising therapeutic target^10–12^. During the late replication phase, the Gag and Gag-Pol proteins—which contain the CA domain essential for protein interactions—are expressed^13^. HIV-1 protease (PR) cleaves the precursor Gag, thereby releasing CA, which then self-assembles into the mature fullerene-cone-shaped mature capsid composed of approximately 250 hexamers and 12 pentamers^14–17^. This structure is essential to the virus for post-entry steps^18,19^. Upon fusing with the host membrane, the viral core enters the cytoplasm and undergoes uncoating. This process is regulated by host factors such as CPSF6 and CypA through interaction with the CA. These interactions are essential for efficient reverse transcription and subsequent integration of the viral genome^20^.

LEN binds to the N-terminal domain (NTD)-CTD interface of CA monomers, located within a hydrophobic FG (phenylalanine-glycine motif)-binding pocket^2^. The site, situated near the intrahexamer interface of the mature capsid is critical for proper uncoating during cytoplasmic transport^21,22^. LEN reportedly accelerates capsid assembly, leading to aberrant capsid formation, and inhibition of infection^3^. Additionally, it destabilizes Gag and Gag-Pol, thereby interfering with precursor Gag assembly and release^1,2^. Although the FG-binding pocket is absent in immature capsids, it remains unclear whether LEN inhibits viral release at this stage. Although previous studies on the inhibitory effects of LEN have primarily focused on viral maturation and post-entry processes^1,2,21,23,24^, its impact on viral release remains poorly understood. Moreover, assessments of viral release have largely relied on p24 ELISA assays^1,5^, limiting the depth of analysis. In this study, we employed mass photometry, velocity gradient centrifugation, and electron microscopy to investigate the effects of LEN on late-stage of HIV-1 replication. Through a comprehensive analysis of released viral particles, we elucidated the mechanism by which LEN inhibits HIV-1 during the late phase of the viral replication process.

## Results

### LEN does not inhibit HIV-1 release

We isolated LEN-resistant HIV-1 from patients treated with LEN and identified three critical amino acid substitutions (referred to as LRM) in the CA domain that confer resistance to LEN (in preparation). To evaluate the inhibitory effect of LEN on viral release, we measured Vpr-HiBiT levels in virions from 293T/Vpr-HiBiT cells. During Gag assembly, Vpr-HiBiT was incorporated into the progeny virions. Nanoluciferase activity measured via HiBiT and LgBiT complementation indirectly reflects the number of released viral particles^25^. LEN did not reduce the nanoluciferase activity in either X4- and R5-tropic HIV-1, with an IC_50_ value ˃100 nM (**Figs. 1a and 1b**). Similarly, LEN did not inhibit the release of transmitted/founder HIV-1 clones from transfected 293T/Vpr-HiBiT cells (**Table 1**).

**Figure 1.**
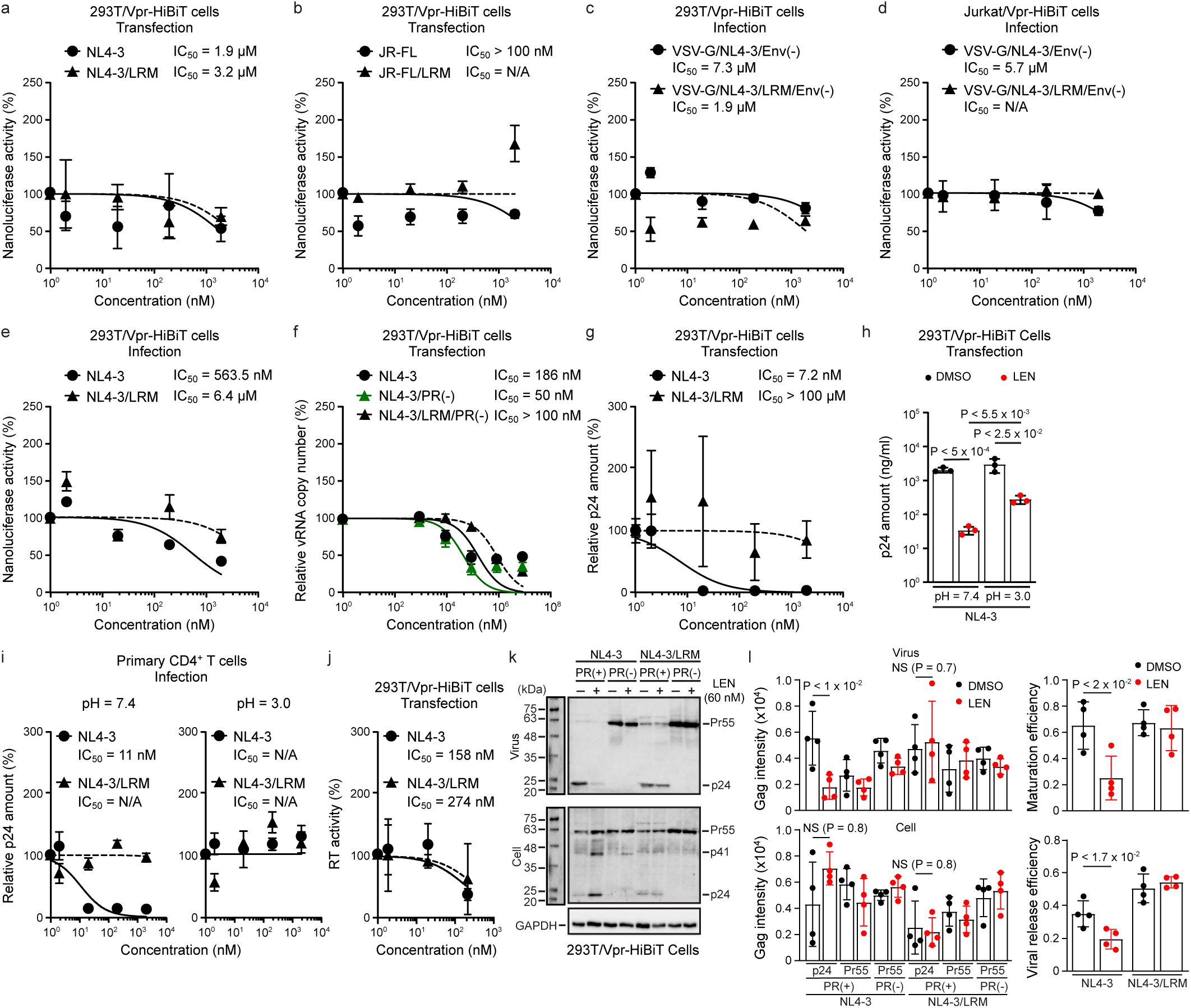
LEN does not inhibit HIV-1 release. **a–e**, Measurement of Vpr-HiBiT amount to assess HIV-1 release from pNL4-3-transfected 293T/Vpr-HiBiT cells (**a**), pJR-FL-transfected 293T/Vpr-HiBiT cells (**b**), VSV-G-pseudotyped NL4-3-infected 293T/Vpr-HiBiT cells (**c**), VSV-G-pseudotyped NL4-3-infected Jurkat/Vpr-HiBiT cells (**d**), and NL4-3-infected Jurkat/Vpr-HiBiT cells (**e**). LEN was added immediately after transfection (**a**,**b**) or 15 h after infection (**c**–**e**). **f**, Measurement of vRNA levels in virions from transfected 293T cells treated with LEN. **g**–**i**, Quantification of p24 Gag in virions from LEN-treated transfected 293T/Vpr-HiBiT cells (**g**,**h**) and primary CD4^+^ T cells (**i**). p24 Gag was measured under acidic conditions (**h**,**i**). **j**, Quantification of reverse transcriptase (RT) activity in virions from NL4-3-transfected 293T/Vpr-HiBiT cells. **k**, Representative western blot showing mature Gag (24 kDa), immature Gag (55 kDa), and GAPDH following LEN treatment (60 nM). **l**, band intensities quantified using ImageJ. Data are presented as mean ± s.d. from two independent (**i**), three independent experiments (**a**– **g**,**h**,**j**), and four independent experiments (**k**,**l**) .

Furthermore, VSV-G-pseudotyped NL4-3 and NL4-3/LRM infected into 293T/Vpr-HiBiT cells or Jurkat/Vpr-HiBiT cells, exhibited no inhibition by LEN when treated 15 h post-infection, with IC_50_ values ˃7.3 μM and 5.7 μM, respectively (**Figs. 1c and 1d**). Replication-competent NL4-3 and NL4-3/LRM also exhibited no substantial inhibition at 15 h post-infection, with IC_50_ values ˃563 nM and 6.4 μM, respectively (**Fig. 1e**). These values were notably higher than previously reported picomolar IC_50_ values^1,5^. However, LEN strongly inhibited infection when added at the time of infection, consistent with previously reported post-entry antiviral effects **(Extended Data** Figs. 1a–1d)^1,5^. LEN also had no effect on vRNA copy numbers, with IC_50_ ˃186 nM (**Fig. 1f**). PF74 and BVM did not block viral release, which is consistent with previous studies^4,7^ **(Extended Data** Figs. 1e and 1f). Overall, these results demonstrate that LEN does not inhibit HIV-1 release during the late stages of replication.

### LEN masks the HIV-1 p24 epitope but is reversed under acidic conditions

To quantify the p24 antigen levels in the supernatants from transfected 293T/Vpr-HiBiT cells and infected-primary CD4^+^ T cells, we used both an in-house and commercial p24 ELISA kits^26^. LEN reduced p24 detection over 100-fold, with an IC_50_ of 7.2 nM and 11 nM in 293T cells and CD4^+^ T cells, respectively (**Figs. 1g** and **1i; Extended Data Fig. 2a**). These values exceed those reported previously^1^, and contradict the Vpr-HiBiT and vRNA measurements (**Figs. 1a–1f**). LEN binds tightly to FG-binding pockets in the CA domain via hydrogen bonds. We hypothesized that this interaction masks the p24 epitopes recognized by the monoclonal antibody. To validate this, 2 μM LEN was added to lysed virions. This resulted in a five-fold decrease in p24 detection **(Extended Data** Fig. 2b), which supported our hypothesis. Subsequently, we explored whether physical treatments could reverse the masking effect. Boiling in SDS buffer failed to restore p24 levels. Notably, freezing samples in PBS enhanced detection 10-fold in the presence of LEN **(Extended Data** Figs. 2c and 2d). As PBS becomes mildly acidic when frozen^27^, the increased acidity may expose otherwise masked epitopes. Histidine protonation is known to alter the CA structure under acidic conditions^28^. Given that one (His62) of the six conserved histidine residues is located adjacent to the LEN-binding site, protonation may expose the FG-binding pocket and restore antibody recognition. As expected, incubating virions under acidic conditions (pH 5.0–3.0) restored p24 detection by up to ten-fold in the presence of LEN (**Figs. 1h and 1i; Extended Data Fig. 2e**). These results indicate that LEN structurally masks the p24 epitope, and this effect can be reversed under acidic conditions, facilitating antibody binding. As masking likely depended on the specific antibody epitopes, previous ELISA-based evaluations of LEN’s effect on viral release may have overestimated its inhibitory activity because of epitope interference.

### HIV-1 Gag-Pol assembly in virions is unaffected by LEN

To assess whether LEN interferes with Gag-Pol packaging, the RT activity in virions produced from transfected cells was measured. LEN treatment did not inhibit the RT activity (**Fig. 1j**). In the absence of PR, Pr55 Gag levels and release of immature particles were unchanged in cells and virus lysates, despite LEN treatment (**Fig. 1k**). Even when using a polyclonal antibody (HIV-Ig) to avoid epitope masking, the p24 Gag level still decreased in conjunction with LEN. Consequently, viral release efficiency, calculated as viral Gag over total Gag, was considerably reduced. This suggests that LEN interferes with antibody detection rather than with actual virion release (**Fig. 1l; Extended Data** Figs. 3a and 3b). Furthermore, LEN did not inhibit the Gag-Pol incorporation into progeny virions. Its apparent effects on proteolytic processing and maturation may be artifactual, resulting from p24 epitope masking.

### LEN alters the HIV-1 Gag distribution

Unlike mature HIV-1 Gag, which contains multiple FG-binding pockets essential for LEN binding, the Pr55 Gag lacks these defined pockets^15,29^. Our data demonstrated that LEN did not reduce Pr55 Gag levels in cell lysates nor did it inhibit the release of immature HIV-1 particles from producer cells following transfection. Notably, p24 levels were higher in cell lysates than in viral lysates (**Fig. 1l**), suggesting the intracellular retention of Gag. Consequently, we hypothesized that LEN binds to Pr55 Gag in the absence of the canonical FG-binding pockets. This was validated by examining the intracellular Gag distribution in the presence of LEN. In both transfected and infected cells, LEN treatment led to a distinct punctate pattern of Pr55 Gag localization, in contrast to the diffuse signals observed under control conditions (**Figs. 2a and 2b**). Notably, LEN had no observable effect on the localization of GagVenus, a Gag protein tagged with yellow fluorescent protein YFP at the C-terminus **(Extended Data** Fig. 4a). Furthermore, supporting super-resolution imaging using direct stochastic optical reconstruction microscopy (dSTORM) revealed significant Gag-Gag aggregation in LEN-treated cells, suggesting enhanced oligomerization of Gag at the plasma membrane (**Figs. 2c and 2d**). This was confirmed via a Gag multimerization assay using a multilayered sucrose gradient to separate Gag monomers from oligomers and higher-order multimers. In cells transfected with the pNL4-3/GagFlag, control samples showed a peak in fraction 4. However, the peak shifted to fraction 5 in LEN-treated samples, indicating the accumulation of heavier oligomeric forms. This shift was not observed in the NL4-3/LRM, which exhibited comparable behavior under both the control and LEN-treated conditions. These results suggest that LEN promotes abnormal Gag oligomerization in the plasma membrane (**Figs. 2e and 2f**) by binding exclusively to the CA-NTD portion, constituting the FG-binding pocket in the mature CA hexamer. The molecular mechanism underlying this abnormal Gag oligomerization was elucidated through molecular simulations, which revealed that LEN binds to immature CA or Pr55 Gag **(Extended Data** Fig. 4b). However, this interaction appears unstable, likely owing to the structural flexibility of immature Gag. Moreover, LEN was predicted to cause steric clashes between residues E79 and R82 at the interface of immature CA monomers, potentially disrupting the hexameric CA-SP1 lattice **(Extended Data** Fig. 4c). This disruption was capable of destabilizing the six-fold symmetry of the lattice **(Extended Data** Fig. 5), altering the particle density and Gag oligomerization. Overall, LEN interacts with immature Gag, destabilizing the CA hexamer structure and promoting abnormal oligomerization at the plasma membrane.

**Figure 2.**
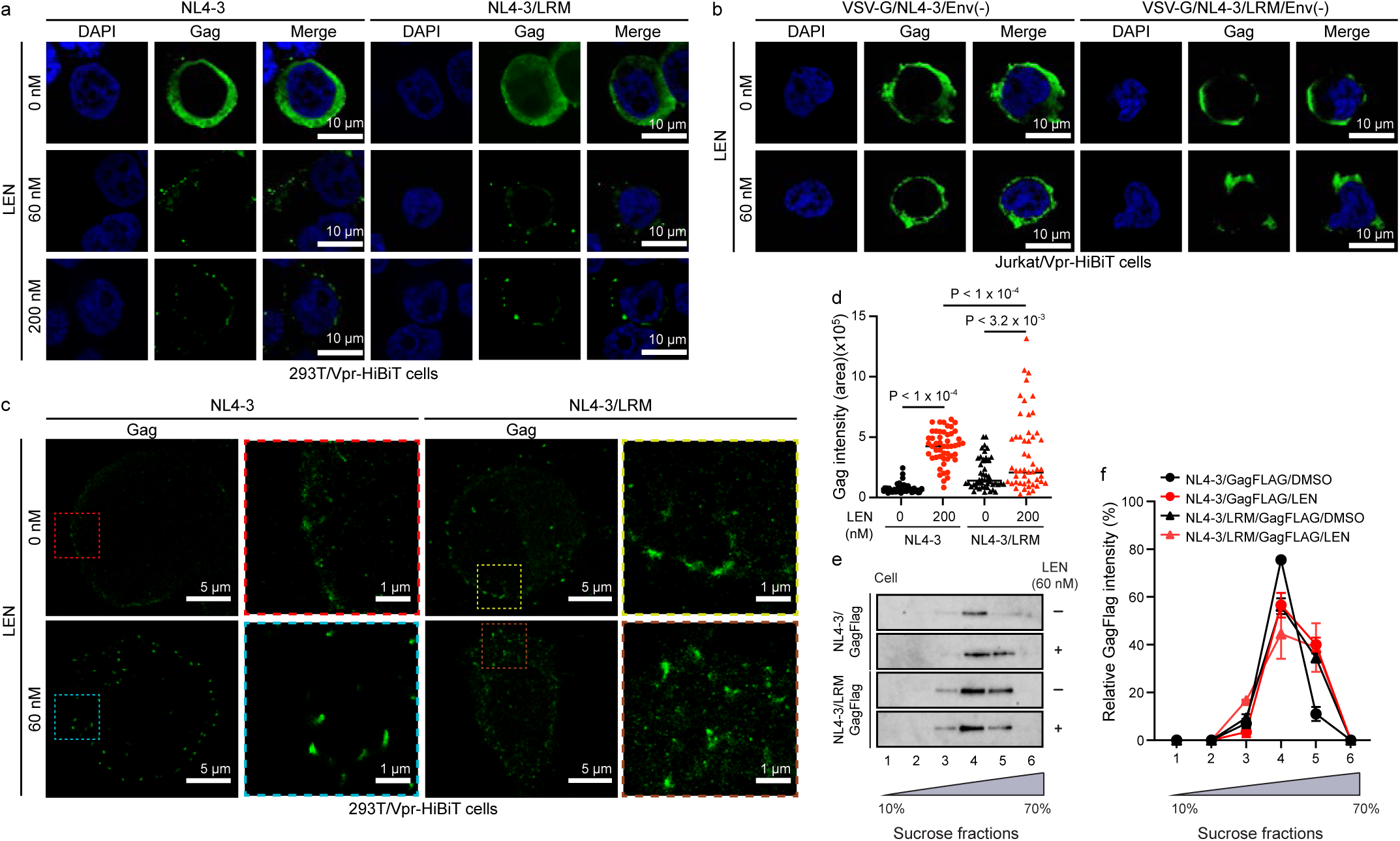
LEN induces the abnormal multimerization of Gag at the plasma membrane. **a**,**b**, Representative confocal images of pNL4-3-transfected 293T/Vpr-HiBiT cells (**a**) or VSV-G/NL4-3-infected Jurkat/Vpr-HiBiT (**b**) treated with LEN. Cells were stained for Gag (green) and nuclei (blue); scale bar, 20 μm. **c**, Representative dSTORM images of pNL4-3-transfected 293T/Vpr-HiBiT cells treated with LEN; scale bars: 5 μm (left panels) and 0.5 μm (enlarged panels). **d**, Gag signal intensities were quantified using ImageJ, with measurements taken from 50 areas across 10 cells. Data are presented as mean ± s.d. from three independent experiments. **e**, Western blotting analysis of Gag multimerization in cells treated with LEN using sucrose gradient analysis. **f**, Band intensities in each fraction were quantified with ImageJ. Data are presented as mean ± s.d. from three independent experiments.

### LEN alters the size and density of HIV-1 particles released from producer cells

During HIV-1 Gag assembly, approximately 1500 to 2000 Gag molecules are assembled at virus-forming regions to produce each progeny virion^30^. We hypothesized that LEN disrupts the orderly assembly of Gag at the plasma membrane, thereby affecting the formation and physical properties of the virus particles. This was validated using a sucrose velocity gradient (5–30%) based on Stokes’ principle, which separates particles based on size and density following ultracentrifugation. Under control conditions, both mature (NL4-3) and immature virions (NL4-3/PR(-)) peaked in fraction #7 **(Extended Data** Fig. 6a). In contrast, treatment with LEN caused a shift in particle distribution, with a peak observed in fraction #11 (**Fig. 3a; Extended Data** Fig. 6a). Additionally, a higher concentration (200 nM) of LEN significantly reduced the number of particles in fraction #7 by two-to three-fold **(Extended Data** Fig. 6b). Contrastingly, a lower concentration (4 nM) had minimal effects **(Extended Data** Fig. 6b). Similarly, 293T/Vpr-HiBiT cells infected with VSV-G-pseudotyped NL4-3 and NL4-3/LRM exhibited identical results **(Extended Data** Fig. 6d). Predictably, PF74, a compound with similar capsid-targeting properties, also induced a comparable shift **(Extended Data** Fig. 6e), whereas BVM did not **(Extended Data** Fig. 6f). This aligns with the findings of Keller *et al*, who reported unaltered particle sizes with BVM, even at high concentrations^31^. The infectivity of the particles separated by the sucrose gradient was evaluated by examining the virions in each fraction. Amongst the untreated samples, fractions #6 and #7 contained infectious viruses. However, in the presence of LEN, these fractions lost their infectivity. Notably, fraction #8 from LEN-treated NL4-3/LRM (60 nM) did not retain infectivity (**Fig. 3b**), suggesting that NL4-3/LRM is resistant at the late stage of viral lifecycle but remains susceptible to loss of infectivity at higher doses of LEN. These findings indicate that LEN significantly alters the physical characteristics of HIV-1 particles, specifically their density and size, likely attributable to increased Gag oligomerization during assembly.

**Figure 3.**
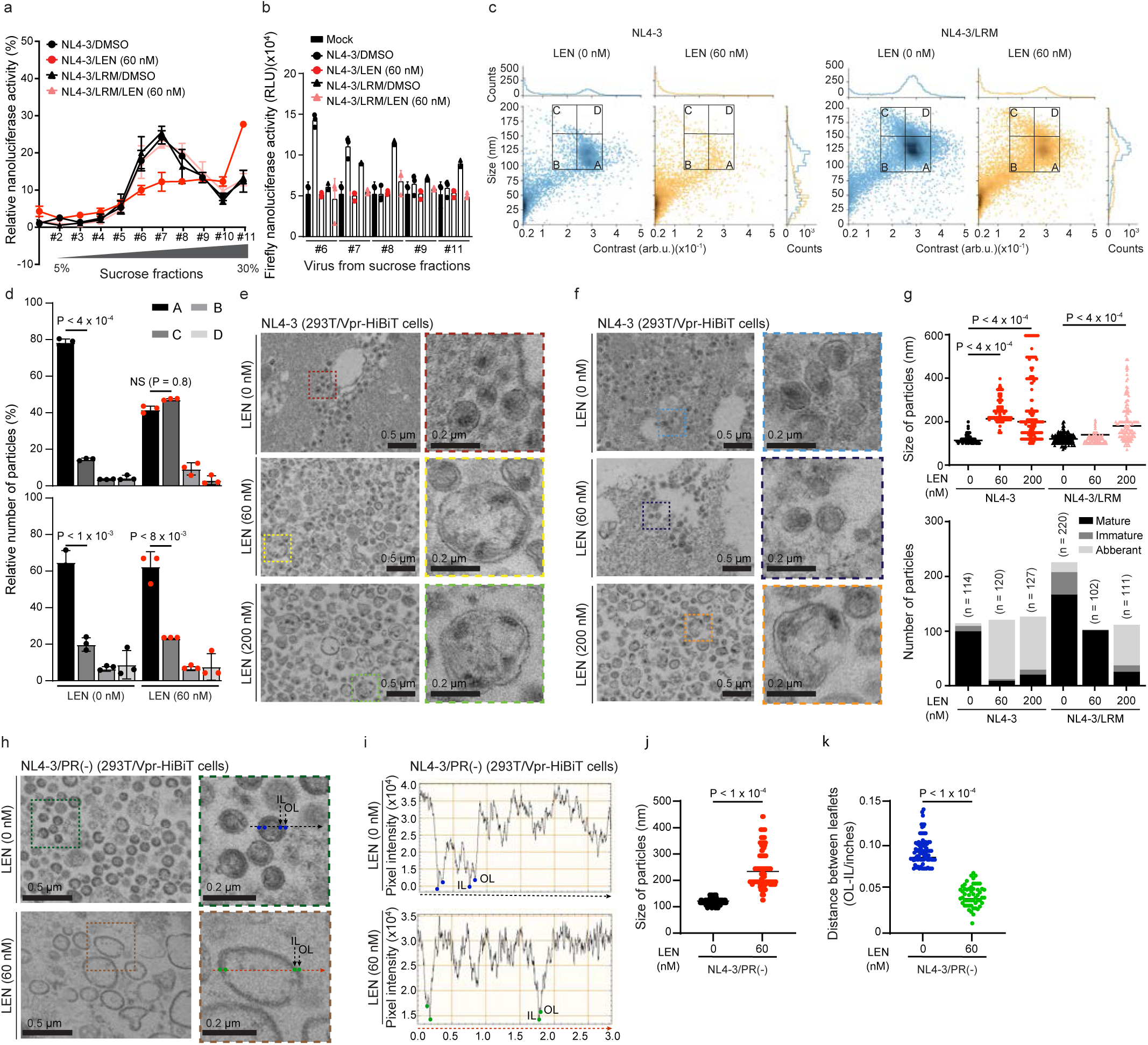
LEN alters the size and density of HIV-1 particles. **a**, Sucrose velocity gradient analysis of HIV-1 particles released from pNL4-3 transfected 293T/Vpr-HiBiT cells treated with LEN, quantified based on Vpr-HiBiT levels. **b**, Infectivity of virions purified from sucrose fractions shown in **a**, assessed by firefly luciferase activity in TZM-bl cells following normalization to Vpr-HiBiT. **c**,**d**, Mass photometry analysis of virion size (nm) and density (contrast; arbitrary units) in the presence (orange) or absence (blue) of LEN. The relative cell number in each gate (A–D) shown in **c** was quantified in **d**. Data are presented as mean ± s.d. from three independent experiments (**a**–**d**). **e**,**f**, Representative transmission electron microscopy (TEM) images of virions produced from pNL4-3-transfected 293T/Vpr-HiBiT cells treated with LEN. Colored dashed boxes indicate enlarged regions. Scale bars, 0.5 μm (main) and 0.2 μm (inset). **g**, Quantification of virion size and classification (mature; black, immature; dark gray, aberrant; light gray) from TEM images in **e** and **f**, performed using ImageJ. *n* indicates the number of virions analyzed. Statistical analysis was performed using one-way ANOVA with Sidak’s correction. **h**, TEM images of NL4-3/PR(-) particles produced from transfected 293T/Vpr–HiBiT cells treated with LEN. Colored dashed boxes indicate enlarged regions. Scale bars, 0.5 μm and 0.2 μm. **i**–**k**, Quantification of Radial density profiles (**i**), particle size (**j**), and outer leaflet (OL)–inner leaflet (IL) distance (**k**) from images in **h**, using ImageJ. Statistical analysis was performed using one-way ANOVA with Sidak’s correction.

### LEN disrupts regular HIV-1 particle size and density

The velocity gradient results were further validated via mass photometry, which utilizes interferometric scattering (iSCAT) detection to assess particle size (≤200 nm) and contrast (density)^32^. Analysis of progeny virions harvested from the producer cells revealed that LEN treatment caused a leftward shift in the particle density distribution of NL4-3 virions. Specifically, there was a considerable increase in the number of particles falling within Gate B and a corresponding decrease in Gate A, where regular-sized viral particles were typically observed (**Fig. 3c and 3d**). LEN also induced a similar shift in particle distribution of immature virions **(Extended Data** Fig. 7a). This reduction in normal-sized particles was especially pronounced at higher LEN concentrations in both NL4-3 and NL4-3/LRM but was not evident at lower LEN concentrations **(Extended Data** Figs. 8a and 8b). Under normal conditions, HIV-1 particles typically measure 100–130 nm in diameter and contain approximately 1500–2000 Gag molecules^30,33^. Mass photometric analysis revealed that these particles displayed a consistent density of approximately 0.30. LEN treatment altered this typical particle profile, yielding a density value of 0.25. Similar changes were observed in VSV-G-pseudotyped NL4-3 virions derived from Jurkat/Vpr-HiBiT cells exposed to increasing concentrations of LEN **(Extended Data** Fig. 7b), as well as in PF74-treated transfected cells **(Extended Data** Fig. 8c). Contrastingly, BVM treatment did not noticeably alter the particle size or density **(Extended Data** Fig. 8d), which was consistent with our sucrose gradient results. Although the reduction in particle number within Gate A did not fully align with the vRNA copy number and nanoluciferase measurements (**Figs. 1a–1f**), these results suggest that LEN disrupts the formation of regular-sized viral particles by altering the particle density and reducing the proportion of mature virions.

### LEN increases HIV-1 particle size

To address the discrepancies between the mass photometry results and Vpr-HiBiT quantification, we used transmission electron microscopy (TEM) to visualize the morphology of HIV-1 particles released from 293T and T cells. Mature HIV-1 particles are characterized by a dark central density, which corresponds to the mature capsid core^34^. Owing to sample sectioning during TEM preparation, capsids may not consistently appear conical or fullerene-shaped; particles can still be reliably classified as mature, immature, or abnormal **(Extended Data** Fig. 9a; **Table 2**). In the presence of LEN, NL4-3 particles exhibit high variability in morphology and size, with many particles exceeding 200 nm in diameter (**Figs. 3e–3g**). This effect was also observed in NL4-3/PR(-) particles (**Figs. 3h–3j**), making them undetectable by mass photometry, as it exceeds its upper size detection limit. A similar particle size heterogeneity was observed in 293T/Vpr-HiBiT cells infected with VSV-G-pseudotyped NL4-3 and NL4-3/LRM **(Extended Data** Figs. 9b, 9c, and **9e).** Scanning electron microscopy (SEM) revealed of heterogeneity among virions attached to the cell surface **(Extended Data** Fig. 9d). These abnormally large, irregular particles are referred to as LEN-induced virus-like particles (LENiVLPs). Notably, in NL4-3/PR(-) particles treated with LEN, LENiVLPs appeared more spherical with a decreased radial distance between the Gag-inner and -outer cell membranes of the viral envelope (**Fig. 3i and 3k**). Overall, these results suggest that LEN interacts with immature Gag and promotes the formation of abnormal large LENiVLPs, likely by altering the structural organization and radial spacing of the viral membrane layers.

### LEN inhibits viral infectivity

LEN inhibits HIV-1 infectivity^1,2^. To assess the infectivity of LENiVLPs, virions were harvested 48 h post-transfection from 293T/Vpr-HiBiT cells, and normalized based on Vpr-HiBiT levels. They were subsequently used to infect TZM-bl, CEM-GFP, and Jurkat cells. Viral infectivity in TZM-bl cells was assessed by measuring HIV-1 Tat-driven firefly luciferase activity at 48 h post-infection. LEN inhibited the infectivity of NL4-3 with an IC_50_ that was 3.4 times lower than that of NL4-3/LRM, indicating increased sensitivity of NL4-3 towards LEN (**Fig. 4a**). Similarly, LEN also inhibited the infectivity of JR-FL-derived virions (**Fig. 4b**). PF74 demonstrated similar infectivity inhibition at 5 μM, consistent with previous reports^4^ **(Extended Data** Fig. 10a). BVM exhibited similar effects **(Extended Data** Fig. 10b). Additionally, when NL4-3 produced from LEN-treated 293T/Vpr-HiBiT cells was used to infect CEM-GFP cells, the IC_50_ was reduced by 8-fold (**Fig. 4c**). In contrast, infection of Jurkat cells resulted in a 3-fold reduction in IC_50_ (**Fig. 4d; Extended Data** Figs. 10c and 10d). Moreover, virions produced from LEN-treated infected-T cells exhibited a 162-fold reduction in infectivity compared to NL4-3/LRM when tested in TZM-bl cells (**Fig. 4e**). A similar pattern was observed in the virions harvested from primary CD4^+^ T cells (**Fig. 4f**), confirming the broad inhibitory effects of LEN on various T cell types. Despite this inhibition, LENiVLPs successfully incorporated envelope proteins during the late-stage assembly in the presence of LEN (**Figs. 4g and 4h Extended Data Fig. 10e**). These particles maintained an effective membrane fusion capacity (**Fig. 4i**), suggesting that LEN inhibits infectivity at the post-entry step, specifically by blocking viral uncoating and/or the nuclear import of reverse transcription complexes.

**Figure 4.**
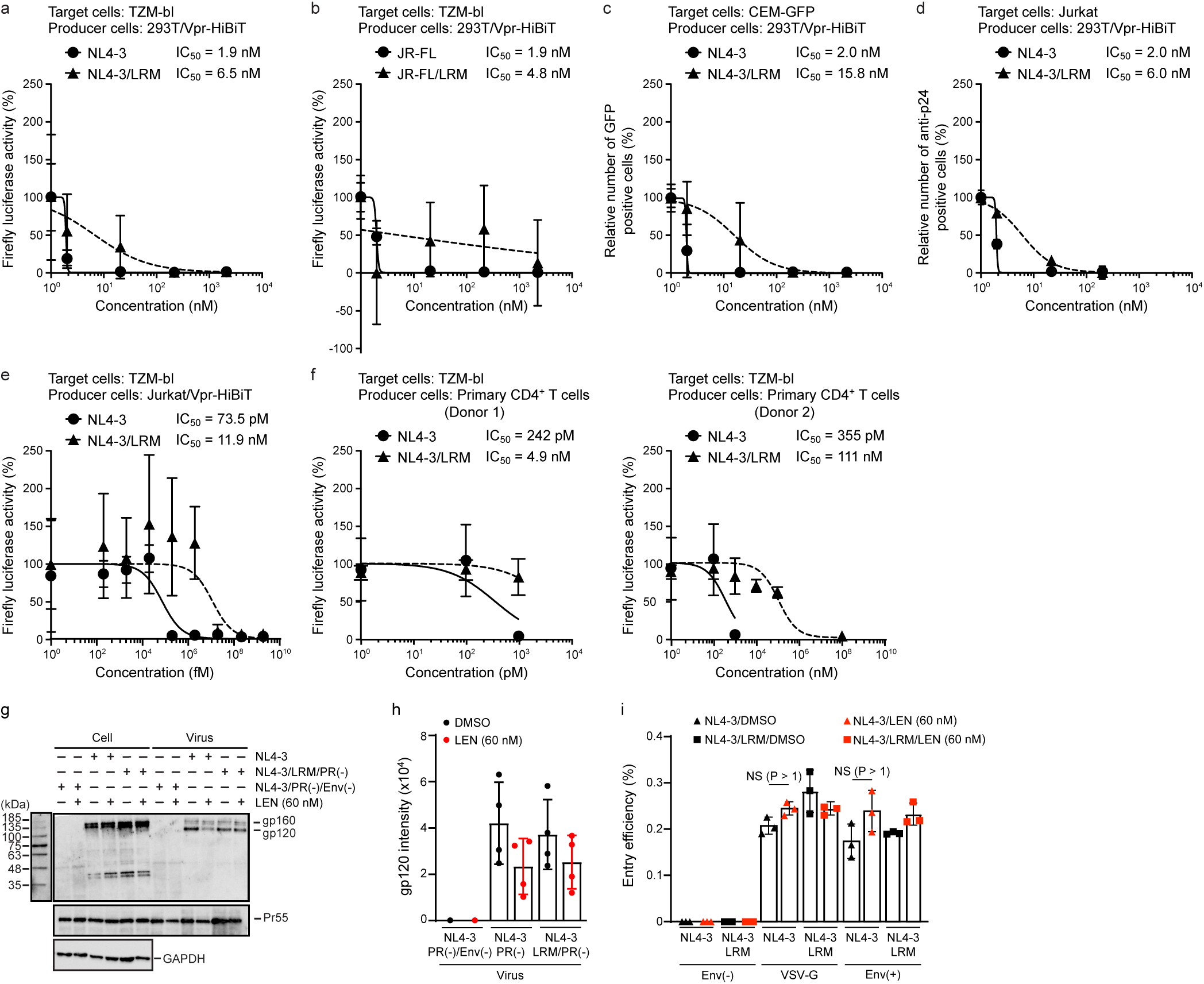
LEN inhibits HIV-1 viral infectivity at a post-entry step. **a**,**b**, Infectivity of HIV-1 virions produced from pNL4-3-transfected (**a**) and pJR-FL-transfected (**b**) 293T/Vpr-HiBiT cells treated with LEN, assessed by Tat-driven firefly luciferase activity in TZM-bl cells. **c**, Infectivity of virions from pNL4-3-transfected 293T/Vpr-HiBiT cells measured in CEM-GFP cells by flow cytometry, based on GFP-positive cells. **d**, Infectivity of virions from pNL4-3-transfected 293T/Vpr-HiBiT cells measured in Jurkat cells by flow cytometry, based on intracellular p24-staining. **e**,**f**, Infectivity of HIV-1 virions produced from pNL4-3-infected Jurkat/Vpr-HiBiT cells (**e**) and pNL4-3-infected primary CD4^+^ T cells (**f**) treated with LEN, assessed by Tat-driven firefly luciferase activity in TZM-bl cells. Data are presented as mean ± s.d. from three independent experiments (biological triplicates where applicable) (**a**–**f**). **g**, Representative western blot showing HIV-1 gp120 incorporation in the presence or absence of LEN. **h**, Quantification of gp120 incorporation normalized to p24 levels, based on ImageJ analysis. Data are presented as mean ± s.d. from four independent experiments. **i**, HIV-1 fusion assessed using a BlaM-Vpr reporter assay and flow cytometric detection of CCF4 dye cleavage in the presence of LEN. Data are presented as mean ± s.d. from three independent experiments.

## Discussion

LEN, a potent capsid inhibitor, binds to the FG-binding pockets located at the mature interfaces between p24 capsid subunits^2^, thereby inhibiting nuclear import, virion assembly, and core formation^1,2^. However, its role in the late-stage of HIV-1 life cycle remains poorly understood. Our findings provided important insights into the effects of LEN on viral assembly, release, and infection. However, these effects were diminished in LEN-resistant virus (LRM) isolated from HIV-1-infected individuals. Notably, LEN failed to inhibit viral release in both 293T and T cells, in contrast to earlier reports^1,2^. This discrepancy is highlighted by our data demonstrating that HIV-1 vRNA copy number, RT activity, and Vpr-HiBiT release remained unaffected by LEN, even at higher concentrations, whereas p24 levels were significantly reduced. Presumably, LEN masks key epitopes on the capsid, thereby interfering with the binding of the p24 antibody. Epitope masking has been reported to occur due to ligand binding that persists even after SDS-induced denaturation^35^, and cannot be ruled out in the case of LEN. Evidence for epitope masking was validated when freezing of lysed samples restored detectable p24 antigen levels, whereas boiling had no such effect^27^.

Furthermore, this supports previous findings that freezing can alter the pH, affecting the HIV-1 capsid structure^28^. These structural changes, in turn, could expose specific regions (epitopes) that influence antibody detection. Additionally, protonation of histidine residues may trigger structural transitions. The p24 capsid contains six histidine residues (His12, His62, His84, His87, His120, and His226). Protonation of these residues may induce electrostatic repulsion, destabilize the interface, and expose epitopes masked by LEN, thereby facilitating antibody recognition. Notably, masking may not occur if the antibody target regions are far from the LEN-binding site.

LEN stabilizes the hexameric lattices of mature HIV-1 capsids via interaction with the FG-binding pockets, but may not bind immature capsids lacking these pockets^2^. However, high-resolution *in situ* cryo-EM has demonstrated that LEN binds to immature Gag despite the absence of FG-binding pockets^29^, which is consistent with simulation data indicating disruption at the CA-SP1 interface (**Extended Data** Figs. 4b and 4c). In this study, LEN had no effect on total Pr55 Gag levels in either cells or viral lysates, suggesting no effect on immature virion production. Nonetheless, the reduced p24 signal in the viral lysates, likely due to epitope masking, makes it unclear whether maturation was impaired. Confocal and dSTORM imaging revealed enlarged Gag puncta with LEN, indicating that LEN alters Gag distribution and promotes its aggregation at the plasma membrane. This supports the ability of LEN to interact with immature Gag and induce oligomerization even in the absence of FG-binding pockets. With regard to GagVenus, large puncta were present at the plasma membrane even without LEN, and their size remained unchanged upon treatment **(Extended Data** Fig. 4a). It is also possible that LEN binds less efficiently to GagVenus, warranting further investigation.

LENiVLPs appear to incorporate Env proteins during late-stage assembly. The incorporated Envs remained functional and mediated efficient membrane fusion (**Figs. 4g–4i**). Env is highly mobile and forms prominent clusters on mature particles, which are essential for efficient target cell binding^36^. This indicates that Gag within LENiVLPs is cleaved by proteases. Despite their enlarged size and aberrant cores, LENiVLPs efficiently attach to and fuse with target cells, suggesting that particle size and core morphology do not substantially affect these early post-entry steps. These findings reinforce the inference that LEN inhibits the post-entry stage of the viral replication cycle.

We also identified giant viral particles, termed LENiVLPs, with aberrant capsid structures in both the transfected and infected cells. We propose that LEN disrupts Gag-Gag interactions at assembly sites, resulting in abnormal Gag accumulation at the plasma membrane and the formation of these enlarged particles. Destabilization of the CA-SP1 hexameric structure by LEN likely impairs hexamer-hexamer interactions, further promoting Gag oligomerization and altering particle density (**Fig. 3c; Extended Data** Figs. 4c and 5). We observed a reduced radial distance between the Gag layer (appearing as inner leaflet) and the plasma membrane (appearing as outer leaflet) in LEN-treated samples (**Figs. 3k**). This could be attributed to the structural collapse or modification of the MA domain, which anchors Gag to the plasma membrane. Alternatively, Gag multimers may adopt a flatter conformation, leading to decreased spacing between outer and inner leaflets. Additional structural studies, such as cryo-EM, are necessary for conclusively determining this. While performing cryo-EM, mature LENiVLPs should not be dismissed as a rare species, given their diverse and abnormal morphologies.

Overall, these results expand our current understanding of LEN’s antiviral mechanism by demonstrating its ability to target both mature and immature Gag assemblies, as revealed by advanced virus analysis techniques that incorporate cutting-edge technologies. This dual-phase inhibition—through the disruption of precursor Gag assembly and proper uncoating— presents new opportunities for refining capsid-targeting therapeutics.

## Acknowledgments

This work was supported by the Japan Agency for Medical Research and Development (AMED) Research Program on HIV/AIDS (JP25fk0410075h0001, JP25fk0410058h0003, JP24fk041047h0003, and JP25fk0410056h0003) to A.S.; AMED Research Program on HIV/AIDS (JP24fk0410058h0001) to Y.I.; AMED (JP21fk0410026h0001, JP23fk0410058h0001, JP24fk0410065h0001, and JP24fk0410058h0202) to K.M., AMED (JP24fk0410058h0001) to Y.I. JSPS KAKENHI Grants (JP23KK0292 and JP23K06561) to K.M.; the JSPS KAKENHI Grant-in-Aid for Scientific Research (C) JP24K09227 to A.S.; the JSPS KAKENHI Grant-in-Aid for Scientific Research (B) JP22H02500 to A.S. and JP21H02361 (to A.S.); JSPS KAKENHI Grant-in-Aid for Scientific Research (C) 22K07103 to T.I.; the JSPS Bilateral Program JPJSBP120245706 to A.S.; the JSPS Fund for the Promotion of Joint International Research (International Leading Research) JP23K20041 (to A.S.); the G-7 Grants (2024) to A.S.; Shionogi infectious disease research promotion foundation (2024) to A.S.; the Ito Foundation Research Grant R6 KEN119 (to A.S.);, and the

Takeda Science Foundation to K.M. and T.I; the Mitsubishi Foundation to T.I.; International Joint Research Project of the Institute of Medical Science, the University of Tokyo to T.I. The following reagent was obtained through the NIH HIV Reagent Program, Division of AIDS, NIAID, NIH: polyclonal Anti-Human Immunodeficiency Virus Immune Globulin, Pooled Inactivated Human Sera, ARP-3957, contributed by NABI and National Heart Lung and Blood Institute (Dr. Luiz Barbosa), Lenacapavir, HRP-20266, contributed by the NIH HIV Reagent Program, Zidovudine (AZT) (#HRP-3485) and anti-HIV-1 p24 Gag Monoclonal (#24-4), ARP-6521, contributed by Dr. Michael Malim, CEM-GFP cells, ARP-3655, contributed by Dr. Jacques Corbeil, and TZM-bl cells, HRP-8129, and pMM310 (#ARP-11444), contributed by Dr. Michael Miller (Merck Research Laboratories). pJR-FL was provided by Dr. Yoshio Koyanagi. p24 ELISA kit was kindly provided from Dr. Yuetsu Tanaka and Dr. Reiko Tanaka. Dr. Shinichiro Nishimura and Dr. Motohide Takahashi assisted with the initial electron microscopy imaging.

## Author’s Contributions

KM, YI, TS, YM conceived and coordinated the study. WAOA conducted majority of the experiments in this study and prepared the original manuscript. PN, MJH, and KM provided support for plasmid construction and data analysis. HT performed mass photometry analysis. HO conducted a structural analysis using computer simulations. HS and TM provided technical support for electron microscopy. NM prepared the electron microscopy samples and conducted the imaging. TI provided the RT assay technique and membrane fusion analysis technique using the BlaM assay. AS assisted with the experiments involving transmitted/founder viruses. YN performed quantification of vRNA. All the authors have read and approved the final manuscript.

## Materials and Methods

### Ethics statement

The study protocol was approved by the Institutional Review Board for General Studies at the Faculty of Life Sciences, Kumamoto University. Written informed consent was obtained from all participants in accordance with the Declaration of Helsinki.

### Chemical compounds

LEN (SUNLENCA; formerly GS-6207) synthesized at Gilead Sciences (Foster City, CA, USA) was obtained from MedChem Express (Cat. No. HY-111964; New Jersey, USA) and via the NIH AIDS Research and Preference Reagent Program, Division of AIDS at the National Institute of Allergy and Infectious Diseases. Puromycin was purchased from Sigma-Aldrich (Cat. No. ant-pr-1; Massachusetts, USA). PF74 was purchased from MedChem Express (Cat. No. HY-120072; New Jersey, USA). BVM was purchased from SelleckChem (Cat. No. S9010; Houston, Texas, USA, specifically at 9330 Kirby drive, STE 200, Houston, TX 77054). Zidovudine (AZT; #HRP-3485) manufactured by TCI America, Inc, (Oregon, United States) was sourced from NIH AIDS Research and Preference Reagent Program, Division of AIDS at the National Institute of Allergy and Infectious Diseases.

### Plasmids

For pJR-FL and pNL4-3, mutations of Q67H, K70R, and T107N in the p24 capsid region were introduced by site-directed mutagenesis. The D25N protease-inactivating mutation was inserted into pNL4-3 to generate pNL4-3/PR(-) and pNL4-3/LRM/PR(-). The pRDI292/Vpr-HiBiT construct was created by tagging the C-terminus of Vpr with an 11-amino-acid HiBiT via PCR using primers containing *BamHI* and *SalI* sites^25^. Specifically, the first and second forward primers shared the sequence 5’- GTATCGGATCCATGGAACAAGCCCCAGAAGACCAA-3’, which encoded the *vpr* sequence and included a *BamHI* restriction site. The first reverse primer (5’- CGGCTGGCGGCTGTTCAAGAAGATTAGCTAGGTCGACCAGCTGTG-3’) encoded the HiBiT tag and a *SalI* site, while the second reverse primer (5’- GAAATGGAGCCAGTAGATCCGTGAGCGGCTGGCGGCTGTTCAAGA-3’) encoded the HiBiT tag fused to HIV-1 Vpr. The resulting PCR product (Vpr-HiBiT) was inserted into pRDI292 vector^37^ using *BamHI* and *SalI* restriction enzymes. To reconstruct the intact 5′-long terminal repeat (LTR), the U3 region was inserted upstream of the pRDI292/Vpr-HiBiT transcription start site. The final construct, pRDI292/Vpr-HiBiT encodes an LTR-driven Vpr-HiBiT fusion protein and SV40-driven puromycin resistance genes. To prevent packaging of the Vpr-HiBiT genes into HIV-1 virions, the packaging signal was removed from the pRDI292/Vpr-HiBiT construct.

pNL4-3/Env(-) was generated by deleting a segment of the env gene between the two *BglII* restriction sites (nucleotides 7032 to 7612) within the *gp120* coding region, resulting in a frameshift mutation that prevents the expression of functional gp160 Env protein^38^. To construct pNL4-3/GagFLAG, a two-step PCR was performed. The first forward primer included a *BssHII* site (5’- CTCGCCTCTTGCCGTGCGCGCTTCAGCAAGCCGAGTCCTGC-3’), and the first reverse primer encoded the FLAG tag (5’- TTACTTGTCGTCATCGTCTTTATAATCCCCGGGTTGTGACGAGGGGTCGCTGC-3’). A second PCR was performed using the first PCR product as a template, with the same forward primer and a reverse primer encoding *SbfI* site (5’- TTTTTCTGTTTTAACCCTGCAGGTTACTTGTCGTCATCGTCTTT-3’). PCR products were purified and digested with *DpnI* for 1 h, followed by enzyme inactivation. The pNL4-3 vector was digested with *BssHII* and *SbfI*, gel-purified, and ligated with the PCR products using NEBuilder HiFi DNA Assembly Master Mix (Cat. No. E2621X; New England Biolabs, Massachusetts, USA). The ligation mixture was transformed into *E. coli* strain JM109 cells and incubate overnight. For the BlaM fusion assay, pCXN/NLEnv and pMM310, which encodes BlaM-Vpr, were used as described previously^39^.

### Cells

TZM-bl cells (also known as JC.53.bl-13), a HeLa derived cell line that stably expresses high levels of CD4 and CCR5 receptors^40^, were obtained through the AIDS Research and Reference Reagent Program, Division of AIDS, NIAID, NIH, from Dr. John C. Kappes, Dr. Xiaoyun Wu and Tranzyme Inc^41^. TZM-bl cells harbor Tat-driven reporter genes encoding firefly luciferase (Luc) and *Escherichia coli* β-galactosidase. TZM-bl, HEK293T, 293T/Vpr-HiBiT, and HeLa cells were cultured in Dulbecco’s modified Eagle medium (Cat. No. D5796; DMEM; Sigma-Aldrich, Missouri, USA) supplemented with 5% fetal bovine serum (FBS; Cat. No. 173012; Sigma-Aldrich). Jurkat, Jurkat/Vpr-HiBiT, and CEM-GFP cells were maintained in RPMI 1640 medium (Cat. No. 11875; Thermo Fisher Scientific, Massachusetts, USA) with 10% FBS (RPMI-10). CEM-GFP cells, which express green fluorescent protein (GFP) under the control of the HIV-1 LTR promoter, were obtained through the AIDS Research and Reference Reagent Program, Division of AIDS, NIAID, NIH, from Dr. Jacques Corbeil^42^.

Peripheral blood mononuclear cells (PBMCs) were isolated from whole blood by density gradient centrifugation using Ficoll-Paque (Cat. No. 50494; Organon Teknika Co., North Carolina, USA), followed by two washes with PBS containing 2% FBS to remove platelets and contaminants. PBMCs were resuspended at 1 × 10⁷ cells/mL in isolation buffer. To isolate untouched CD4^+^ T cells by negative selection, the PBMC suspension was incubated with a biotinylated antibody cocktail targeting non-CD4^+^ cell surface markers (e.g., CD8, CD14, CD19, and CD56) using the human CD4^+^ T Cell Isolation kits (Cat. No. 130-096-533; Miltenyi Biotec, Tokyo, Japan) for 10-15 min at 4 °C. Streptavidin-coated magnetic beads (Cat. No. 11-161-D; Dynabeads Human T-Activator CD3/CD28, Gibco, Thermo Fisher Scientific) were added to bind the labeled CD4^+^ T cells. The cell mixture was then cultured at 37 °C in RPMI-10 supplemented with 20 ng/mL phytohemagglutinin (PHA; Cat. No. L9132; Sigma-Aldrich) for 3 days. After activation, the suspension was placed in a magnetic separator for 2-5 min, and the supernatant containing CD4^+^ T cells was carefully collected, centrifuged at 300 × g for 5 min, and resuspended in RPMI-10 containing PHA (10 µg/ml).

Jurkat/Vpr-HiBiT and 293T/Vpr-HiBiT cell lines were previously established^25^. To generate these cell lines, the plasmid pRDI292/Vpr-HiBiT was linearized using the restriction enzyme *HpaI*. The linearized plasmid was transfected into HEK/293T cells using Lipofectamine 3000 (Cat. No. L3000015; Thermo Fisher Scientific) according to manufacturer’s instructions. For Jurkat cells, the linearized plasmid was introduced by electroporation using a NEPA21 Type II electroporator (Nepa Gene Co., Ltd. Chiba, Japan). As the packaging signal had been removed from pRDI292/Vpr-HiBiT, lentiviral transduction was not possible. After transfection and electroporation, cells were cultured in puromycin-containing medium to select for stable integrants expressing the puromycin-resistance gene.

### Virus harvesting method

293T/Vpr-HiBiT cells (2 × 10^4^ cells/well in a 12-well plate) were transfected with one of the following plasmids: pNL4-3, pNL4-3/LRM, pJR-FL, and pJR-FL/LRM, using Lipofectamine 3000. Simultaneously, cells were treated with the indicated concentration of LEN, PF74, or BVM. At 48 h post-transfection, supernatants containing virions were collected and centrifuged at 13,200 × g for 1 h at 4 °C. The resulting viral pellets were resuspended in RPMI-10.

To generate VSV-G-pseudotyped HIV-1, HEK/293T cells were co-transfected with either pNL4-3/Env(-) with pVSV-G or pNL4-3/LRM/Env(-) with pVSV-G using Lipofectamine 3000. At 48 h post-transfection, supernatants were harvested, and virions were pelleted by centrifugation (13,200 × g, 4 °C, 1 h), then resuspended in RPMI-10. These pseudotyped viruses were used to infect 293T/Vpr-HiBiT cells (2 × 10^4^ cells/well in a 12-well plate) and Jurkat/Vpr-HiBiT cells (1 × 10^5^ cells/well in a 12-well plate) by spinoculation (1,190 × g, room temperature, 2 h), with virus amounts normalized to p24 levels (measured by ELISA). LEN was added 15 h post-infection. At 72 h post-infection, supernatants were collected, virions were pelleted by centrifugation, and resuspended in RPMI-10.

For the analysis of replication-competent virus, 293T cells (1 × 10^6^ cells/mL) were transfected with pNL4-3 and pNL4-3/LRM. At 48 h post-transfection, supernatants were collected, pelleted by ultracentrifugation (83,500 × g, 4 °C, 1 h), and resuspended in RPMI-10. Viral concentrations were quantified by p24 ELISA. Jurkat/Vpr-HiBiT and primary CD4^+^ T cells were infected with NL4-3 and NL4-3/LRM by spinoculation. After spinoculation, the unbound virus was removed by washing the cells three times with PBS. LEN and AZT were added 15 h post-infection. Cells were then incubated in RPMI-10 at 37 °C for 4 days. For CD4^+^ T Cells, RPMI-10 was supplemented with PHA (10 µg/ml).

### Virus quantification

The amount of released Vpr-HiBiT (indicating viral particle release) was quantified by measuring nanoluciferase activity using the Nano-Glo® HiBiT Lytic Detection System (Cat. No. N3040; Promega, Wisconsin, USA). Briefly, culture supernatants were lysed with Nano-Glo® HiBiT Lytic Buffer, followed by the addition of LgBiT protein and substrate.

Luminescence was then measured using a GloMax® Discover System (GM3000; Promega). The copy number of vRNA in the supernatants was determined by RT–digital PCR (RT-dPCR) using a QIAcuity Nanoplate 26K 24-well instrument and a QIAcuity OneStep Advanced Probe Kit on a QIAcuity dPCR system (Cat. No. 250132; Qiagen, Venlo, The Netherlands). vRNA was extracted from the culture supernatants using a QIAamp Viral RNA Mini Kit (Cat. No. 52904; Qiagen) and eluted with RNase-free distilled water. DNase treatment was applied to the columns during the extraction process. Each vRNA sample was then subjected to RT-dPCR analysis using the following primer and probe set: forward, 5’- AGCCTCAATAAAGCTTGCCTTGA-3’; reverse, 5’- CCCTGTTCGGGCGCCACTGCTAGAG-3’; and probe, 5’-6-FAM (FAM, Carboxyfluorescein) TCTGGTAACTAGAGATCCCTCAGACC-TAMRA (TAMRA, Carboxytetramethylrhodamine)-3’. The probe targets the poly-A signal loop region within the HIV-1 LTR region. RT-dPCR reactions were carried out with the following conditions: RT extension (50 °C for 40 min), initial denaturation (95 °C for 2 min), following by 40 cycles of denaturation (95 °C for 5 s) and annealing-extension (60 °C for 30 s).

The qPCR-base product-enhanced RT (PERT)-assay was performed as previously described^43^. Briefly, culture supernatants were mixed with RNA lysis buffer [0.25% Triton X-100, 50 mM KCl, 100 mM Tris-HCl (pH 7.4), 40% glycerol, 0.8 U/μl recombinant RNase inhibitor (Cat. No. 2313A; Takara Bio Inc., Shizuoka, Japan)] and incubated at room temperature for 10 min. The resulting lysate was used for analysis by real-time PCR following manufacturer’s instructions using the KAPA SYBR Fast qPCR Kit (ROX Low qPCR) (Cat. No. KK4602; NIPPON Genetics, Tokyo, Japan) with the following primers: forward, 5’- TCCTGCTCAACTTCCTGTCGAG-3’ and reverse, 5’- CACAGGTCAAACCTCCTAGGAATG-3’. Fluorescence was measured using the ABI 7300 Real-Time PCR System (Thermo Fisher Scientific).

The amount of p24 capsid protein in the supernatant was quantified according to the manufacturer’s instructions using both a previously described in-house p24 ELISA kit^26^ and a commercially available kit (Cat. No. 0801008; ZeptoMetrix, Buffalo, USA). For heat treatment, SDS sample buffer was added to the pelleted virions, followed by boiling at 95 °C for 10 min. The amount of p24 was then measured by ELISA. For the freeze-thaw experiments, lysis buffer was added to the samples, followed by a 100-fold dilution with PBS and storage at -20 °C overnight. After freezing, p24 concentrations were determined by ELISA.

### Viral infectivity assay

To evaluate viral infectivity, viruses harvested from LEN-treated 293T/Vpr-HiBiT cells were first normalized based on Vpr-HiBiT (nanoluciferase activity), and subsequently used to infect CEM-GFP cells. At 2 days post-infection, the cells were fixed with 4% paraformaldehyde (PFA) (Cat. No. 163-20145; Fujifilm, Osaka, Japan). In parallel, Jurkat cells were also infected with Vpr-HiBiT-normalized virus and fixed with 4% PFA at 2 days post-infection. Following the permeabilization with 0.2% Saponin (S4521; Sigma-Aldrich), the cells were stained with an anti-p24-FITC monoclonal antibody (Cat. No. 6604665; KC57-FITC; Beckman Coulter Inc, California, USA). The percentage of GFP-positive and p24-positive cells was quantified by flow cytometry using FACSCalibur system (BD Biosciences, California, USA).

TZM-bl cells were also infected with virus normalized by Vpr-HiBiT activity. In addition, viruses harvested from LEN-treated CD4^+^ T cells and Jurkat/Vpr-HiBiT cells, normalized by p24 levels, were used to infect TZM-bl cells. The p24 levels were measured under acidic conditions (pH 3). Two days post-infection, firefly luciferase activity in TZM-bl cells was measured using a luciferase assay system according to the manufacturer’s instruction (Promega).

### Sucrose velocity gradient analysis

Sucrose velocity gradient analysis was performed as previously described^44^. Cell-free supernatants containing virions were first centrifuged at 8,000 × g for 5 min to remove cellular debris. The virions were then pelleted by centrifugation at 13,200 × g for 1 h at 4 °C and resuspended in 1 mL of RPMI-10. Each concentrated viral sample was layered onto a 5–30% sucrose gradient (10 mL total volume) and subjected to ultracentrifugation at 83,500 × *g* for 1 h at 4 °C using an SW41Ti rotor in a Beckman L-70 ultracentrifuge (Beckman Coulter Inc, USA). After centrifugation, 1 mL fractions were collected sequentially from the top of the gradient. The amount of Vpr-HiBiT in each fraction was quantified using the Nano-Glo® HiBiT Lytic Detection System.

According to Stoke’s law, the sedimentation rate of a virus is influenced by its density and size.

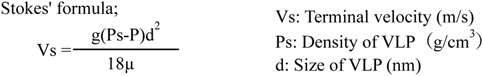

To analyze viral infectivity using a sucrose velocity gradient, virus-containing fractions were first collected. Each 1mL sucrose fraction was diluted with 9 mL of PBS, and the viruses were pelleted by ultracentrifugation at 83,500 × *g* for 1 h at 4 °C. After centrifugation, the purified viruses were quantified using the Vpr-HiBiT assay and subsequently used to infect TZM-bl cells. Two days post-infection, firefly luciferase activity in the TZM-bl cells was measured according to the manufacturer’s instructions (Promega).

### Western blotting analysis

Cells and viruses were lysed with 1% Triton X lysis buffer (1% Triton X-100; Cat. No. 35501-02; Nacalai Tesque, Inc., Kyoto, Japan, 50 mM Tris-HCl, 300 mM NaCl, 10 mM iodoacetamide) supplemented with a protease inhibitor cocktail (Cat. No. 04693116001; Sigma-Aldrich). After lysis, SDS sample buffer was added. HIV-1 Gag, Env, and FLAG-tagged proteins were detected by immunoblotting using the following primary antibodies: HIV-1 Immunoglobulin G (HIV-Ig: NIH AIDS reagent program), anti-HIV-1 gp120 antibody (Cat. No. D7324; Aalto Bio Reagents Ltd. Dublin, Ireland), Anti-DYKDDDDK tag monoclonal antibody (Cat. No. 012-22384; Fujifilm), and anti-p24 Gag monoclonal (#24-4: NIH AIDS Research and Reference Reagent Program). Horseradish peroxidase (HRP)-conjugated anti-mouse (Cat. No. 115-035-003; Jackson ImmunoResearch Laboratories, Pennsylvania, USA) or anti-human secondary antibody (Cat. No. 109-035-003; Jackson ImmunoResearch Laboratories, Pennsylvania, USA) was used. For detection of gp120 protein, HRP-conjugated anti-goat IgG antibody (Cat. No. 305-035-003; Jackson ImmunoResearch Laboratories) was used. HRP signals were visualized using the ChemiDoc Touch Imaging System (BIO-RAD, California, USA).

### Gag multimerization Assay

293T/Vpr-HiBiT cells (4 × 10^5^ cells/well in a 6-well plate) were transfected with either pNL4-3/GagFLAG or pNL4-3/LRM/GagFLAG using Lipofectamine 3000 following the manufacturer’s instructions. Immediately after transfection, cells were treated with the indicated concentrations of LEN. Gag multimerization was analyzed as previously described^45^.

### Viral fusion assay

The HIV-1-based BlaM-Vpr assay was performed with minor modifications as previously described^46^. HIV-1 particles incorporating BlaM-Vpr were generated by co-transfecting HEK/293T cells with pNL4-3/Env(-), pCXN/NLEnv, and pMM310 (encoding BlaM-Vpr). For the negative control, cells were transfected with pNL4-3/Env(-) with pMM310. As a positive control, cells were transfected with pNL4-3/Env(-), pVSV-G, pMM310, with or without LEN treatment. Viruses were harvested 2 days post-transfection. TZM-bl cells (1 × 10^5^ cells/mL), seeded overnight in a 12-well plate, were spinoculated. After spinoculation, the virus-containing medium was removed and replaced with fresh medium. Cells were then incubated at 37 °C for 3 h to allow viral entry. Following incubation, cells were washed three times with PBS and loaded with 2 μM of the cell-permeable fluorescent β-lactamase substrate (CCF4-AM) using the LiveBLAzer FRET-B/G loading kit (Cat. No. K1095; Thermo Fisher Scientific), supplemented with 0.5% solution D (an anion transport inhibitor), in Opti-MEM with 10% FBS. Cells were washed three times with PBS and fixed in 2% PFA. Fluorescence signals at 520 nm (Pacific Blue; uncleaved CCF4) and 447 nm (AmCyan; cleaved CCF4) were measured using a CytoFLEX Flow Cytometer (Beckman Coulter) and analyzed by FlowJo software (v10.7.1, BD Biosciences). Entry efficiency was represented as the ratio of cleaved CCF4 signal to the total signal (cleaved + uncleaved CCF4).

### Transmission electron microscopy analysis

293T/VprHiBiT cells were transfected with pNL4-3, pNL4-3/LRM, and pNL4-3/PR(-) in the presence of LEN. Viruses were collected by centrifugation at 13,200 × *g* for 1 h. After centrifugation, the supernatant was removed, leaving the viral pellet.

VSV-G/NL4-3 harvested from HEK/293T cells was infected into Jurkat/Vpr-HiBiT cells by spinoculation, followed by washing to remove unbound virus. At 15 h post-infection, LEN was added, and the cells were incubated at 37 °C for 4 days. Viruses were then pelleted using the same centrifugation protocol as above.

The pelleted viruses were fixed with 2% glutaraldehyde (Cat. No. G010; GA; TAAB Laboratories Equipment, Aldermaston, England) and 1% osmium tetroxide (Cat. No. O018; TAAB Laboratories Equipment), followed by dehydration through a graded ethanol series (50%, 70%, 80%, 90%, 95%, and 99.5%). Samples were then embedded in Epon812 resin (Cat. No. T024; TAAB Laboratories Equipment). Ultrathin sections were prepared on copper grids (Cat. No. 2823; Nisshin EM, Tokyo, Japan), stained with Mayer’s hematoxylin solution (Cat. No. MHS16; Sigma-Aldrich) and lead citrate (Cat. No. 18-0875-2; Sigma-Aldrich), as described previously^47^. Stained sections were examined using a Hitachi 7600 transmission electron microscope (Hitachi High-Technologies, Tokyo, Japan) at 80 kV.

### Scanning electron microscopy

HeLa cells were seeded onto collagen-coated coverslips. After seeding, the cells were transfected with pNL4-3 plasmid in the presence or absence of LEN. At 48 h post-transfection, the cells on the coverslips were fixed with 2% GA and 1% osmium tetroxide, then dehydrated through a graded ethanol series. The samples were treated with hexamethyldisilazane (Cat. No. 440191; Sigma-Aldrich), air dried, and coated with a thin layer of platinum using an ion sputter. Scanning electron microscopy images were acquired using a JEOL JSM-7200F scanning electron microscope (JEOL, Tokyo, Japan) at 5 kV.

### Confocal/dSTORM microscopy

293T/Vpr-HiBiT were seeded onto collagen-coated 8-well chamber slides (Cat. No. 192-008; Watson Co., Ltd. Tokyo, Japan). Cells were transfected with pNL4-3, pNL4-3/LRM, pNL4-3/GagVenus, or pNL4-3/LRM/GagVenus using Lipofectamine 3000, as described above, and LEN was added immediately after transfection. At 16 h post-transfection, cells were fixed with 4% PFA for 30 min at 4 °C.

For Jurkat/Vpr-HiBiT cells, VSV-G-pseudotyped NL4-3 and NL4-3/LRM were produced in HEK/293T cells, harvested, and concentrated. Jurkat/Vpr-HiBiT cells were infected by spinoculation. After infection, the excess virus was washed off twice with PBS, and the cells were incubated at 37°C for 3 days. LEN was added 15 h post-infection. After incubation, cells were fixed with 4% PFA for 30 min at 4 °C. Permeabilization was carried out using 0.1% Triton X-100 for 2 min for 293T/Vpr-HiBiT cells, and 0.2% Saponin for 2 min for Jurkat/Vpr-HiBiT cells. Both cell lines were then treated with 0.1 M glycine for 10 min, followed by blocking with 3% bovine serum albumin (BSA; Cat. No. 01863-48; Nacalai Tesque Inc.) for 30 min. Then, cells were incubated with a mouse anti-p24 capsid mouse monoclonal antibody (#24-4) for 1 h at 4 °C. After washing, cells were incubated with AlexaFluor 488-conjugated anti-mouse secondary antibody (Cat. No. A-11001; Thermo Fisher Scientific) for 1 h at 4 °C. Nuclei were stained with 4’,6-diamidino-2-phenylindole (DAPI) for 5 min. Samples were then mounted with Dako Fluorescence Mounting Medium (Cat. No. S3023; Dako, Glostrup, Denmark), and imaged using Zeiss LSM 700 laser-scanning confocal microscopy.

For dSTORM analysis, AlexaFluor 546-conjugated anti-mouse secondary antibody (Cat. No. A-21123; Thermo Fisher Scientific) was used, and imaging was performed using the Nanoimager system (Oxford Nanoimaging Limited, Oxford, United Kingdom). Gag-Gag interactions were analyzed using ImageJ by measuring Gag cluster intensities. A standardized area across clusters was used for the measurements.

### Mass photometry

At 48 h post-transfection of 293T/Vpr-HiBiT cells with pNL4-3, pNL4-3/PR(-), pNL4-3/LRM/PR(-), or pNL4-3/LRM in the presence of LEN, PF74, or BVM, the virus-containing supernatant was harvested. The virus was pelleted by centrifugation at 13,200 × *g* for 1 h. For dose-response analysis, VSV-G-pseudotyped NL4-3 and NL4-3/LRM were infected into 293T/Vpr-HiBiT and Jurkat/Vpr-HiBiT cells, followed by the addition of increasing concentrations of drugs at 15 h post-infection. After 4 days, supernatants were collected, and viruses were pelleted using the same centrifugation protocol described above. Virus pellets were resuspended in 2% GA and incubated for 1 h at 4 °C. Samples were then diluted 1:10 with PBS and placed on the MassGlassKV (Cat. No. MGKV; Refeyn Ltd. Oxford, United Kingdom) Subsequently, half the volume (5 µL) was removed and replace with 5 µL of PBS. The samples were then analyzed by mass photometry (Karitro^MP^; Refeyn).

### Computational molecular dynamics simulation

Hexamer models of CA-SP1 with or without six LEN molecules were constructed by homology modeling with Modeller 9v8^48^ based on previously reported structures (PDB 7ASL^49^, 6V2F^1^, 6BHR^50^. In each model, one IP6 molecule was also included. The models were surrounded by water solvent molecules and were then energy-minimized with the steepest descent method, followed by the conjugated gradient method. After heating process up to 310 K for 0.1-ns molecular dynamics (MD) simulations, 25-ns MD simulations at 310 K were conducted. Simulation of CA-SP1 with LEN was duplicated. For the simulations, we used AMBER16 software package (https://ambermd.org/)^51^. We adopted force fields of GAFF for LEN and IP6 and FF14 for the other molecules. Exceptionally, other angle parameters of phosphate^52^ were used for IP6. The time step of 2 fs was applied for the MD simulations.

### Statistical analysis

All statistical analyses were performed using GraphPad Prism9. One-way ANOVA with Sidak’s multiple comparison test and two-way ANOVA with Tukey’s multiple comparisons test were used as appropriate. Paired and unpaired student’s *t*-tests were also performed where indicated. Error bars represent the mean ± standard deviation from at least three independent experiments.

---

**Extended Data Fig. 1: LEN does not inhibit HIV-1 release. a–d**, Antiviral activity of LEN on HIV-1 particle release from infected and transduced cells by measuring the Vpr-HiBiT and p24 amount. **e**,**f**, Antiviral activity of PF74 and BVM on HIV-1 particle release from transfected 293T/Vpr-HiBiT cells by measuring the Vpr-HiBiT amount. Data are mean ± s.d. from three independent experiments.

**Extended Data Fig. 2: HIV-1 release is markedly inhibited by LEN by p24 ELISA. a**, Antiviral activity of LEN against HIV-1 (NL4-3 and JR-FL) release by measuring the p24 antigen amount in the supernatant. Data are mean ± s.d. from three independent experiments. *Two-way ANOVA with Sidak’s multiple comparisons test; p =< 0.05 was considered significant.* **b**, Epitope masking was confirmed by directly adding LEN (6 μM) to harvested virions. The p24 antigen amount in supernatant was determined by p24 ELISA. Data are mean ± s.d. from three independent experiments. *Two-way ANOVA with Sidak’s multiple comparisons test; p =< 0.05 was considered significant*. **c**,**d**, After the addition of p24 lysis buffer, samples were boiled at 95^0^C, after which the p24 antigen amount was analyzed by ELISA. For freezing assay, after the addition of p24 lysis buffer, the samples were diluted with 1xPBS and frozen at -20^0^C. The p24 antigen amount was analyzed by ELISA. Data are mean ± s.d. from three independent experiments *Two-way ANOVA with Sidak’s multiple comparisons test p =< 0.05 was considered significant*. **e**, Lysed samples were diluted with PBS (pH-7.4 to pH-2.0). The p24 amount was measured using p24 ELISA. Data are mean ± s.d. from three independent experiments.

**Extended Data Fig. 3: Antiviral effects of LEN on Gag processing. a,b**, Representative western blots (*uncropped*) experiment with 60 nM LEN treatment showing the antiviral effects of LEN on mature and immature Gag proteins. The band intensities of Gag were measured and analysed using ImageJ software. *Centre line and error bars represents mean ± s.d. values obtained from three independent experiments

**Extended Data Fig. 4 and 5: LEN binding changes the stability of CA-SP1. hexamerization. a**, Representative confocal microscopy images of transfected 293T/Vpr-HiBiT cells in the presence or absence of LEN (60 nM and 200 nM) with pNL4-3/GagVenus. HIV-1/GagVenus (Green) and Nuclei (Blue) images are from three independent experiments, each producing similar results. GagVenus intensity was analysed using ImageJ. Scale bar, 10 μm. **b**, Homology models generated based on PDB 6V2F and 7ASL. Single LEN molecule was forced to place on immature CA hexamer model, by superposing mature CA monomer with LEN into immature CA monomer. **c**, Molecular simulations showing the effect of LEN binding toward CA-SP1 hexamerization. Homology modeling was based on PDB 7ASL, 6V2F, 6BHR with Modeller 9v8. Force field: FF14+GAFF.

**Extended Data Fig. 6: LEN changes the particle size and density of particles. a**, Sucrose velocity gradient of HIV-1/PR(-) particles produced from transfected 293T/Vpr-HiBiT cells in the presence of LEN by measuring the Vpr-HiBiT amount. Data are mean ± s.d. from three independent experiments. **b**,**c**, Sucrose velocity gradient of HIV-1 particles produced from transfected 293T/Vpr-HiBiT cells in the presence of LEN (200 nM and 4 nM) by measuring the Vpr-HiBiT amount. Data are mean ± s.d. from three independent experiments. **d**, Sucrose velocity gradient of HIV-1 particles produced from VSV-G/NL4-3-infected 293T/Vpr-HiBiT cells in the presence of LEN by measuring the Vpr-HiBiT amount. Data are mean ± s.d. from three independent experiments. **e**, Sucrose velocity gradient of HIV-1 particles produced from transfected 293T/Vpr-HiBiT cells in the presence of PF74 (0.2 μM and 5 μM) by measuring the Vpr-HiBiT amount. Data are mean ± s.d. from three independent experiments. **f**, Sucrose velocity gradient of HIV-1 particles produced from transfected 293T/Vpr-HiBiT cells in the presence of BVM (2 nM and 250 nM) by measuring the Vpr-HiBiT amount. Data are mean ± s.d. from three independent experiments.

**Extended Data Fig. 7: Regular size of particles released from producer cell is lost in the presence of LEN. a,** Mass photometry data showing the size (nm) and contrast (density) (arb.u.) of NL4-3/PR(-) virions produced from transfected 293T/Vpr-HiBiT cells in the presence or absence of LEN (60 nM). Data are mean ± s.d. from three independent experiments. **b,** Mass photometry data showing the size (nm) and contrast (density) (arb.u.) of VSV-G-pseudotyped NL4-3/Env(-) and VSV-G-pseudotyped NL4-3/LRM/Env(-) virions produced from infected Jurkat/Vpr-HiBiT cells in the absence (blue) or presence of LEN (2 μM, 200 nM, 20 nM, and 2 nM). Data are mean ± s.d. from three independent experiments.

**Extended Data Fig. 8: Regular size of particles released from producer cell is lost in the presence of PF74 but not BVM. a**,**b**, Mass photometry data showing the size (nm) and contrast (density) (arb.u.) of NL4-3 and NL4-3 LRM virions produced from transfected 293T/Vpr-HiBiT cells in the presence or absence (blue) of LEN (4 nM and 200 nM). Data are mean ± s.d. from three independent experiments. **c**,**d**, Mass photometry data showing the size (nm) and contrast (density) (arb.u.) of NL4-3 and NL4-3/LRM virions produced from transfected 293T/Vpr-HiBiT cells in the absence or presence of PF74 and BVM in increasing concentrations respectively. Data are mean ± s.d. from three independent experiments.

**Extended Data Fig. 9: LEN affect the size of virions produced from 293T and T cells. a,** Classification of virions produced from transfected 293T/Vpr-HiBiT cells. Mature, Immature and abnormal particles was used as classification. Description of these classes are in *Table 2*. **b**,**d**, Representative transmission electron micrograph images and sizes of HIV-1 produced from VSV-G/NL4-3-infected 293T/Vpr-HiBiT and Jurkat/Vpr-HiBiT cells in the presence of LEN. Colored dashed lines represent zoomed sections. Scale bar in Zoomed sections, 0.2 μm. **c**, Size of virus particles was analyzed with ImageJ software **e**, Representative scanning electron micrograph images of HIV-1 produced from transfected HeLa cells in the presence of LEN. Colored dashed lines represent zoomed sections. Scale bar, 0.1 μm.

**Extended Data Fig. 10: Antiviral activity of LEN, PF74 and BVM on viral infectivity. a,b**, Antiviral activity of PF74 and BVM on HIV-1 infectivity of virions produced from transfected 293T/Vpr-HiBiT cells and infected into TZM-bl cells. It was analyzed by measuring the HIV-1 Tat-driven firefly luciferase activity. Data are mean ± s.d. from three independent experiments. **c**, Effect of LEN on viral infectivity of virions released from 293T/Vpr-HiBiT cells and infected into Jurkat cells. The Gag positive cells were analysed by FACS. Data are mean ± s.d. from three independent experiments. **d**, Effect of LEN on viral infectivity of virions released from 293T/Vpr-HiBiT cells and infected into CEM-GFP cells. Viral infectivity was analyzed using flow cytometry by counting the number of GFP-positive cells. Data are mean ± s.d. from three independent experiments. **e,** Quantification of envelope incorporation normalized by p24 by measuring the band intensities using ImageJ software. Data are mean ± s.d. from six independent experiments.

## References

1 Link, J. O. et al. Clinical targeting of HIV capsid protein with a long-acting small molecule. Nature 584, 614–618 (2020). 10.1038/s41586-020-2443-1

2 Bester, S. M. et al. Structural and mechanistic bases for a potent HIV-1 capsid inhibitor. Science 370, 360–364 (2020). 10.1126/science.abb4808

3 Faysal, K. M. R. et al. Pharmacologic hyperstabilisation of the HIV-1 capsid lattice induces capsid failure. Elife 13 (2024). 10.7554/eLife.83605

4 Blair, W. S. et al. HIV Capsid is a Tractable Target for Small Molecule Therapeutic Intervention. Plos Pathog 6 (2010). ARTNe100122010.1371/journal.ppat.1001220

5 Yant, S. R. et al. A highly potent long-acting small-molecule HIV-1 capsid inhibitor with efficacy in a humanized mouse model. Nat Med 25, 1377–1384 (2019). 10.1038/s41591-019-0560-x

6 Fujioka, T. et al. Anti-Aids Agents .11. Betulinic Acid and Platanic Acid as Anti-Hiv Principles from Syzigium-Claviflorum, and the Anti-Hiv Activity of Structurally Related Triterpenoids. J Nat Prod 57, 243–247 (1994). DOI 10.1021/np50104a008

7 Li, F. et al. PA-457: A potent HIV inhibitor that disrupts core condensation by targeting a late step in Gag processing. P Natl Acad Sci USA 100, 13555–13560 (2003). 10.1073/pnas.2234683100

8 Van Baelen, K. et al. Susceptibility of Human Immunodeficiency Virus Type 1 to the Maturation Inhibitor Bevirimat Is Modulated by Baseline Polymorphisms in Gag Spacer Peptide 1. Antimicrob Agents Ch 53, 2185–2188 (2009). 10.1128/Aac.01650-08

9 Gupta, S. K. et al. Lenacapavir administered every 26 weeks or daily in combination with oral daily antiretroviral therapy for initial treatment of HIV: a randomised, open-label, active-controlled, phase 2 trial. Lancet Hiv 10, E15–E23 (2023). 10.1016/S2352-3018(22)00291-0

10 Tedbury, P. R. & Freed, E. O. HIV-1 Gag: An Emerging Target for Antiretroviral Therapy. Curr Top Microbiol 389, 171–201 (2015). 10.1007/82_2015_436

11 Sundquist, W. I. & Kräusslich, H. G. HIV-1 Assembly, Budding, and Maturation. Csh Perspect Med 2 (2012). ARTN a00692410.1101/cshperspect.a006924

12 Carnes, S. K., Sheehan, J. H. & Aiken, C. Inhibitors of the HIV-1 capsid, a target of opportunity. Current Opinion in Hiv and Aids 13, 359–365 (2018). 10.1097/Coh.0000000000000472

13 Mervis, R. J. et al. The Gag Gene-Products of Human Immunodeficiency Virus Type-1 - Alignment within the Gag Open Reading Frame, Identification of Posttranslational Modifications, and Evidence for Alternative Gag Precursors. Journal of Virology 62, 3993–4002 (1988). Doi 10.1128/Jvi.62.11.3993-4002.1988

14 Ganser, B. K., Li, S., Klishko, V. Y., Finch, J. T. & Sundquist, W. I. Assembly and analysis of conical models for the HIV-1 core. Science 283, 80–83 (1999). DOI 10.1126/science.283.5398.80

15 Pornillos, O. et al. X-Ray Structures of the Hexameric Building Block of the HIV Capsid. Cell 137, 1282–1292 (2009). 10.1016/j.cell.2009.04.063

16 Heymann, J. B., Butan, C., Winkler, D. C., Craven, R. C. & Steven, A. C. Irregular and semi-regular polyhedral models for Rous sarcoma virus cores. Comput Math Method M 9, 197–210 (2008). 10.1080/17486700802168106

17 Li, S., Hill, C. P., Sundquist, W. I. & Finch, J. T. Image reconstructions of helical assemblies of the HIV-1 CA protein. Nature 407, 409–413 (2000). 10.1038/35030177

18 Hulme, A. E., Perez, O. & Hope, T. J. Complementary assays reveal a relationship between HIV-1 uncoating and reverse transcription. P Natl Acad Sci USA 108, 9975–9980 (2011). 10.1073/pnas.1014522108

19 Campbell, E. M. & Hope, T. J. HIV-1 capsid: the multifaceted key player in HIV-1 infection. Nat Rev Microbiol 13, 471–483 (2015). 10.1038/nrmicro3503

20 Yamashita, M. & Engelman, A. N. Capsid-Dependent Host Factors in HIV-1 Infection. Trends Microbiol 25, 741–755 (2017). 10.1016/j.tim.2017.04.004

21 Selyutina, A. et al. GS-CA1 and lenacapavir stabilize the HIV-1 core and modulate the core interaction with cellular factors. Iscience 25 (2022). ARTN 10359310.1016/j.isci.2021.103593

22 Li, C. L. et al. Lenacapavir disrupts HIV-1 core integrity while stabilizing the capsid lattice. P Natl Acad Sci USA 122 (2025). ARTN e242049712210. 1073/pnas.2420497122

23 Huang, S. W. et al. The primary mechanism for highly potent inhibition of HIV-1 maturation by lenacapavir. Plos Pathog 21, e1012862 (2025). 10.1371/journal.ppat.1012862

24 Sowd, G. A., Shi, J. & Aiken, C. HIV-1 CA Inhibitors Are Antagonized by Inositol Phosphate Stabilization of the Viral Capsid in Cells. Journal of Virology 95 (2021). ARTN e01445-2110.1128/JVI.01445-21

25 Nyame, P. et al. A heterocyclic compound inhibits viral release by inducing cell surface BST2/Tetherin/CD317/HM1.24. J Biol Chem 300, 107701 (2024). 10.1016/j.jbc.2024.107701

26 Tanaka, R. et al. Suppression of CCR5-tropic HIV type 1 infection by OX40 stimulation via enhanced production of beta-chemokines. AIDS Res Hum Retroviruses 26, 1147–1154 (2010). 10.1089/aid.2010.0043

27 Thorat, A. A. & Suryanarayanan, R. Characterization of Phosphate Buffered Saline (PBS) in Frozen State and after Freeze-Drying. Pharm Res 36, 98 (2019). 10.1007/s11095-019-2619-2

28 Ehrlich, L. S., Liu, T., Scarlata, S., Chu, B. & Carter, C. A. HIV-1 capsid protein forms spherical (immature-like) and tubular (mature-like) particles in vitro: structure switching by pH-induced conformational changes. Biophys J 81, 586–594 (2001). 10.1016/S0006-3495(01)75725-6

29 Wu, C. et al. Structural insights into inhibitor mechanisms on immature HIV-1 Gag lattice revealed by high-resolution in situ single-particle cryo-EM. bioRxiv (2024). 10.1101/2024.10.09.617473

30 Briggs, J. A. G. et al. The stoichiometry of Gag protein in HIV-1. Nat Struct Mol Biol 11, 672–675 (2004). 10.1038/nsmb785

31 Keller, P. W., Adamson, C. S., Heymann, J. B., Freed, E. O. & Steven, A. C. HIV-1 Maturation Inhibitor Bevirimat Stabilizes the Immature Gag Lattice. Journal of Virology 85, 1420–1428 (2011). 10.1128/Jvi.01926-10

32 Kukura, P. et al. High-speed nanoscopic tracking of the position and orientation of a single virus. Nat Methods 6, 923–927 (2009). 10.1038/nmeth.1395

33 Gentile, M. et al. Determination of the Size of Hiv Using Adenovirus Type-2 as an Internal Length Marker. J Virol Methods 48, 43–52 (1994). Doi 10.1016/0166-0934(94)90087-6

34 Nakai, M. & Goto, T. Ultrastructure and morphogenesis of human immunodeficiency virus. J Electron Microsc 45, 247–257 (1996). DOI 10.1093/oxfordjournals.jmicro.a023441

35 Zolotarjova, N. I., Hollis, G. F. & Wynn, R. Unusually stable and long-lived ligand-induced conformations of integrins. J Biol Chem 276, 17063–17068 (2001). 10.1074/jbc.M009627200

36 Chojnacki, J. et al. Envelope glycoprotein mobility on HIV-1 particles depends on the virus maturation state. Nat Commun 8, 545 (2017). 10.1038/s41467-017-00515-6

37 Ariumi, Y., Kuroki, M., Maki, M., Ikeda, M., Dansako, H., Wakita, T., et al. . The ESCRT System Is Required for Hepatitis C Virus Production. PLoS ONE, 6(1): e14517 (2011).

38 St Louis, D. C.., et al. Infectious molecular clones with the nonhomologous dimer initiation sequences found in different subtypes of human immunodeficiency virus type 1 can recombine and initiate a spreading infection in vitro. Journal of Virology 72, 3991–3998 (1998).

39 Maeda, Y., Foda, M., Matsushita, S. & Harada, S. Involvement of both the V2 and V3 regions of the CCR5-tropic human immunodeficiency virus type 1 envelope in reduced sensitivity to macrophage inflammatory protein 1α. Journal of Virology 74, 1787–1793 (2000). Doi 10.1128/Jvi.74.4.1787-1793.2000

40 Platt, E. J., Wehrly, K., Kuhmann, S. E., Chesebro, B. & Kabat, D. Effects of CCR5 and CD4 cell surface concentrations on infections by macrophagetropic isolates of human immunodeficiency virus type 1. J Virol 72, 2855–2864 (1998). 10.1128/JVI.72.4.2855-2864.1998

41 Wei, X. et al. Emergence of resistant human immunodeficiency virus type 1 in patients receiving fusion inhibitor (T-20) monotherapy. Antimicrob Agents Chemother 46, 1896–1905 (2002). 10.1128/AAC.46.6.1896-1905.2002

42 Gervaix, A. et al. A new reporter cell line to monitor HIV infection and drug susceptibility in vitro. P Natl Acad Sci USA 94, 4653–4658 (1997). DOI 10.1073/pnas.94.9.4653

43 Vermeire, J. et al. Quantification of Reverse Transcriptase Activity by Real-Time PCR as a Fast and Accurate Method for Titration of HIV, Lenti- and Retroviral Vectors. Plos One 7 (2012). ARTN e50859 10.1371/journal.pone.0050859

44 Monde, K. et al. Molecular mechanisms by which HERV-K Gag interferes with HIV-1 Gag assembly and particle infectivity. Retrovirology 14 (2017). ARTN 27 10.1186/s12977-017-0351-8

45 Waheed, A. A., Ono, A. & Freed, E. O. Methods for the study of HIV-1 assembly. Methods Mol Biol 485, 163–184 (2009). 10.1007/978-1-59745-170-3_12

46 Begum MM, I. K., Takahashi O, Nasser H, Jonathan M, Tokunaga K, Yoshida I, Nagashima M, Sadamasu K, Yoshimura K, The Genotype to Phenotype Japan (G2P-Japan) Consortium, Sato K and Ikeda T. Virological characteristics correlating with SARS-CoV-2 spike protein fusogen. Front. Virol. 4:1353661 (2024). doi: 10.3389/fviro.2024.135366

47 Sasaki, H. et al. Novel electron microscopic staining method using traditional dye, hematoxylin. Sci Rep-Uk 12 (2022). ARTN 7756 10.1038/s41598-022-11523-y

48 Sali, A. & Blundell, T. L. Comparative Protein Modeling by Satisfaction of Spatial Restraints. Journal of Molecular Biology 234, 779–815 (1993). DOI 10.1006/jmbi.1993.1626

49 Mendonça, L. et al. CryoET structures of immature HIV Gag reveal six-helix bundle. Commun Biol 4 (2021). ARTN 481 10.1038/s42003-021-01999-1

50 Dick, R. A. et al. Inositol phosphates are assembly co-factors for HIV-1. Nature 560, 509-+ (2018). 10.1038/s41586-018-0396-4

51 Case, D. A. et al. The Amber biomolecular simulation programs. J Comput Chem 26, 1668–1688 (2005). 10.1002/jcc.20290

52 Steinbrecher, T., Latzer, J. & Case, D. A. Revised AMBER Parameters for Bioorganic Phosphates. J Chem Theory Comput 8, 4405–4412 (2012). 10.1021/ct300613v

