## Supplementary figures and images for "Lenacapavir binding to immature Gag triggers the emergence of giant HIV-1 virions"

### Supplemental Figure 1

Extended data figure 1

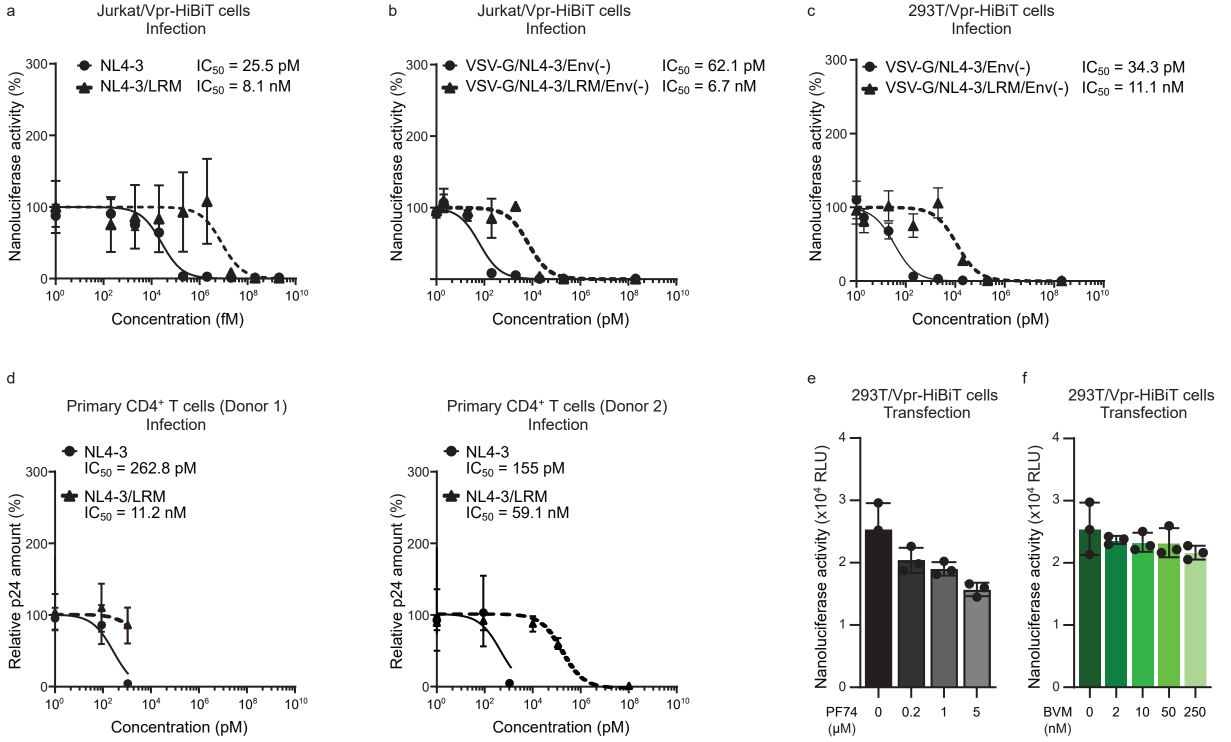

### Supplemental Figure 2

Extended data figure 2

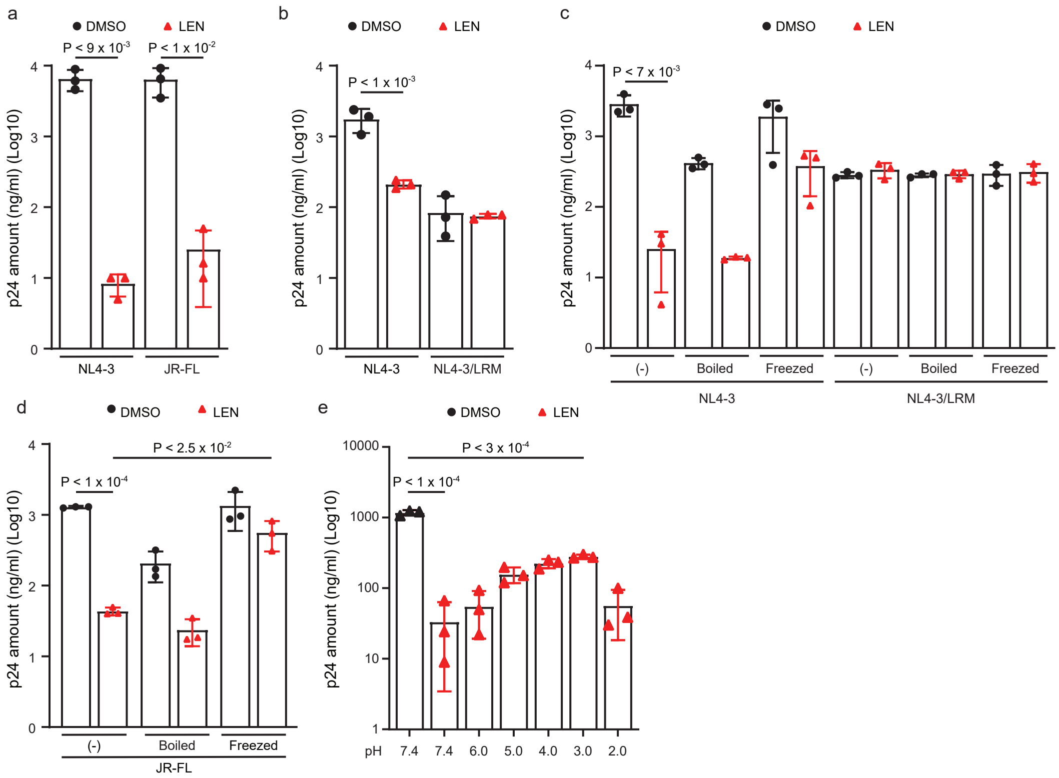

### Supplemental Figure 3

Extended data figure 3

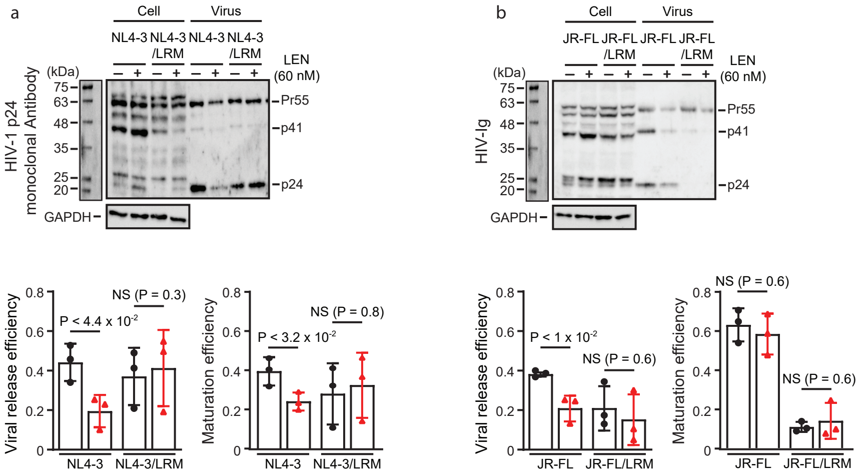

### Supplemental Figure 4

Extended data figure 4

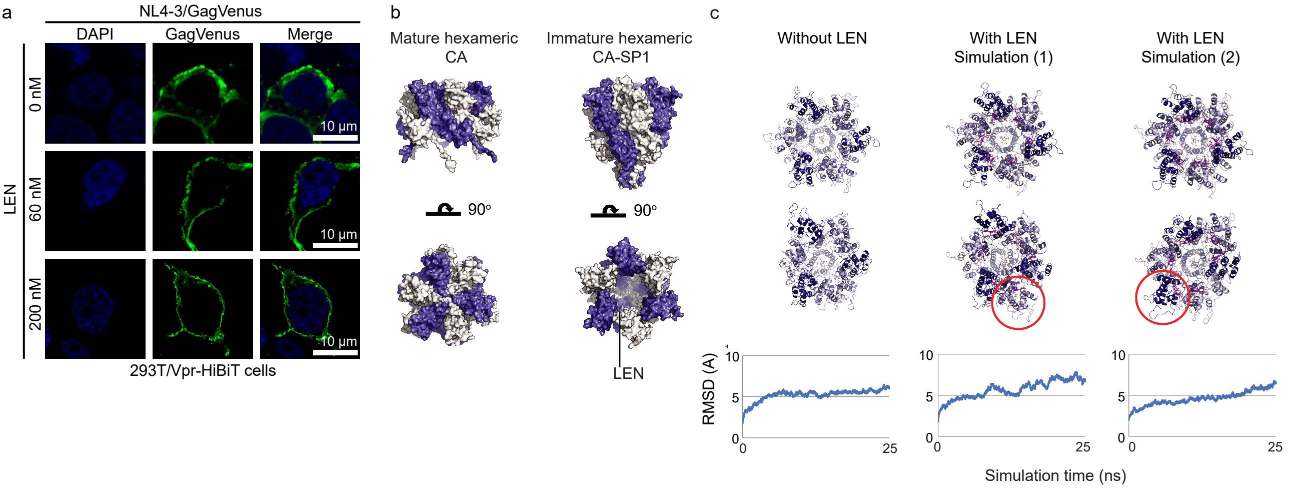

### Supplemental Figure 5

#### Slide 1
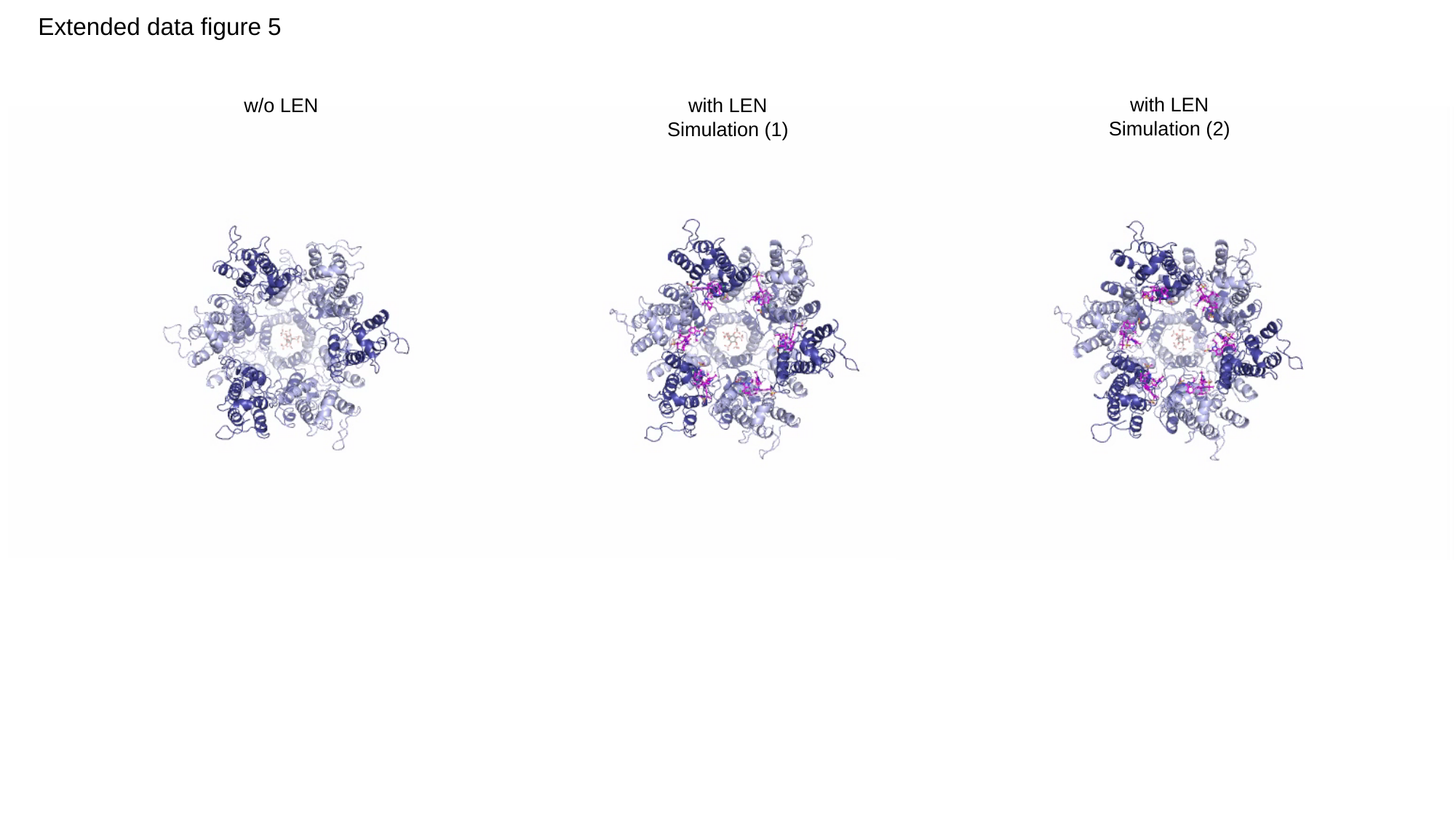

Extended data figure 5
with LEN
Simulation (2)
w/o LEN
with LEN
Simulation (1)

### Supplemental Figure 6

Extended data figure 6

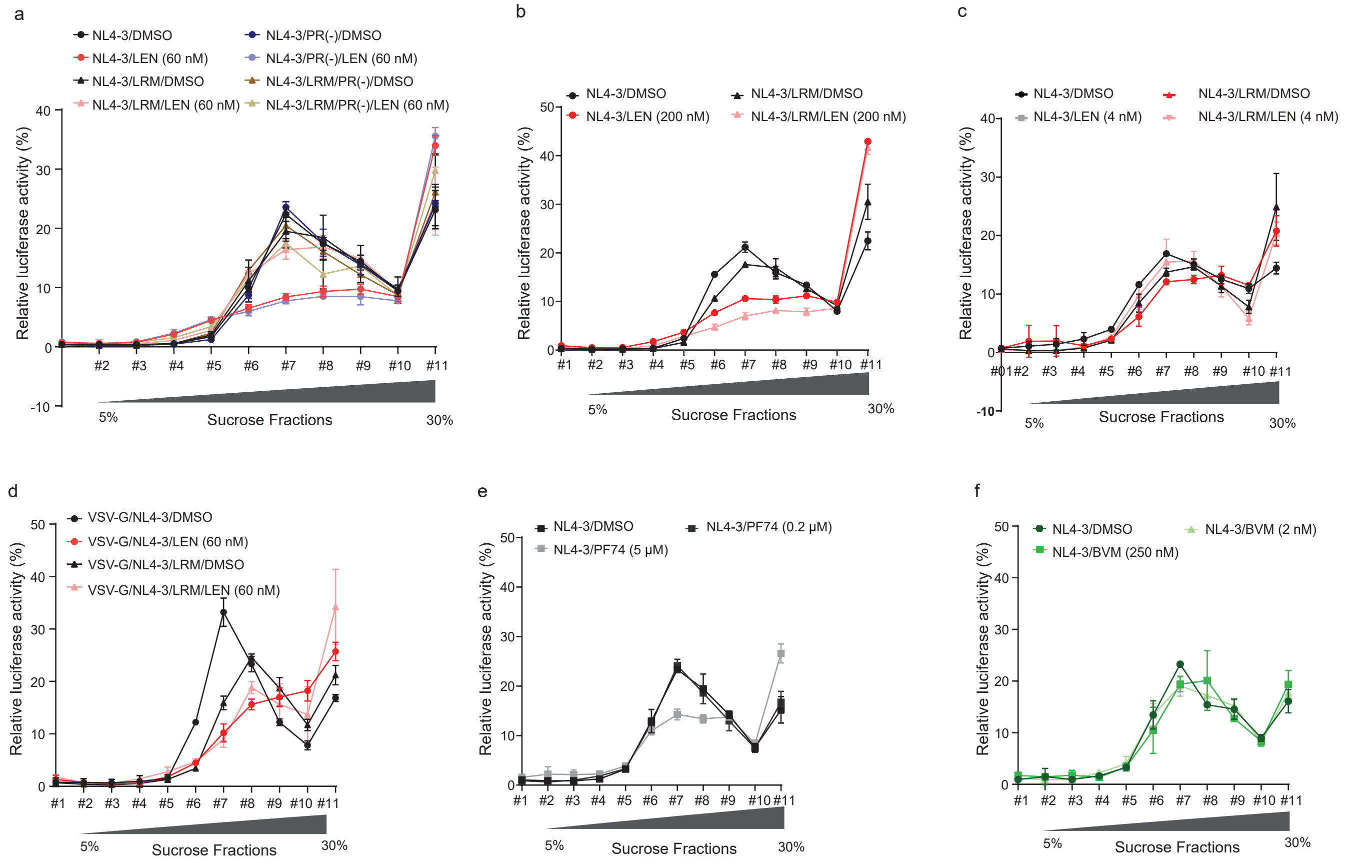

### Supplemental Figure 7

Extended data figure 7

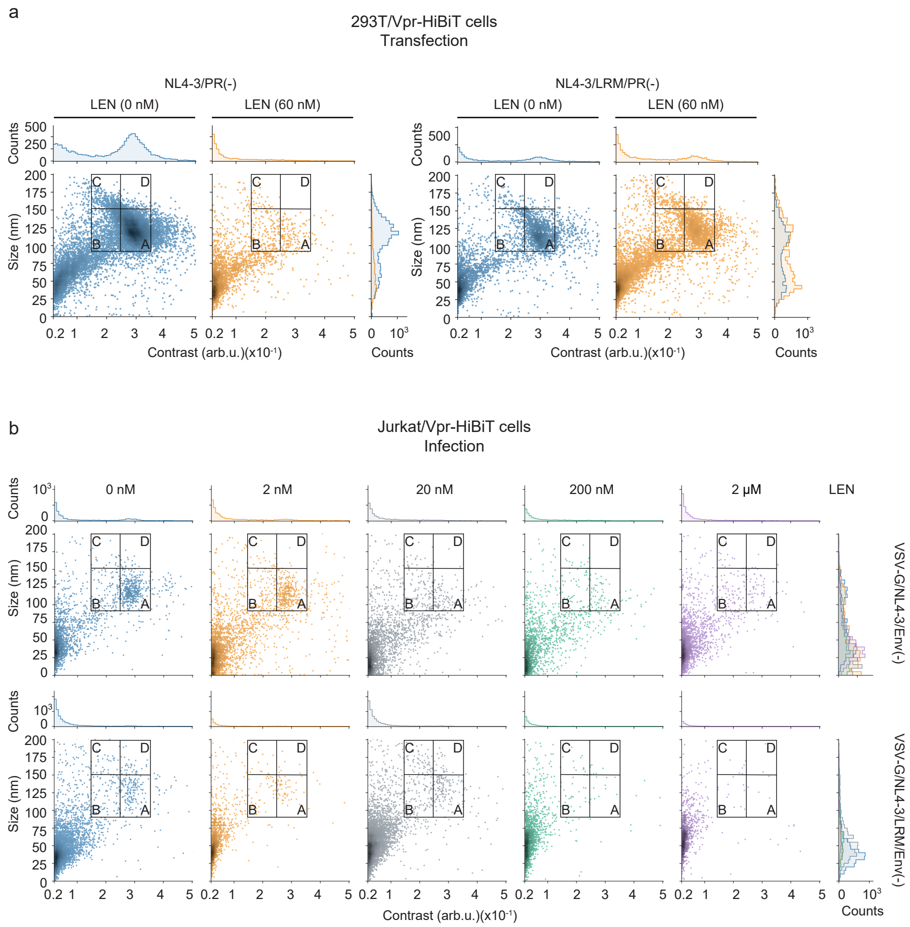

### Supplemental Figure 8

Extended data figure 8

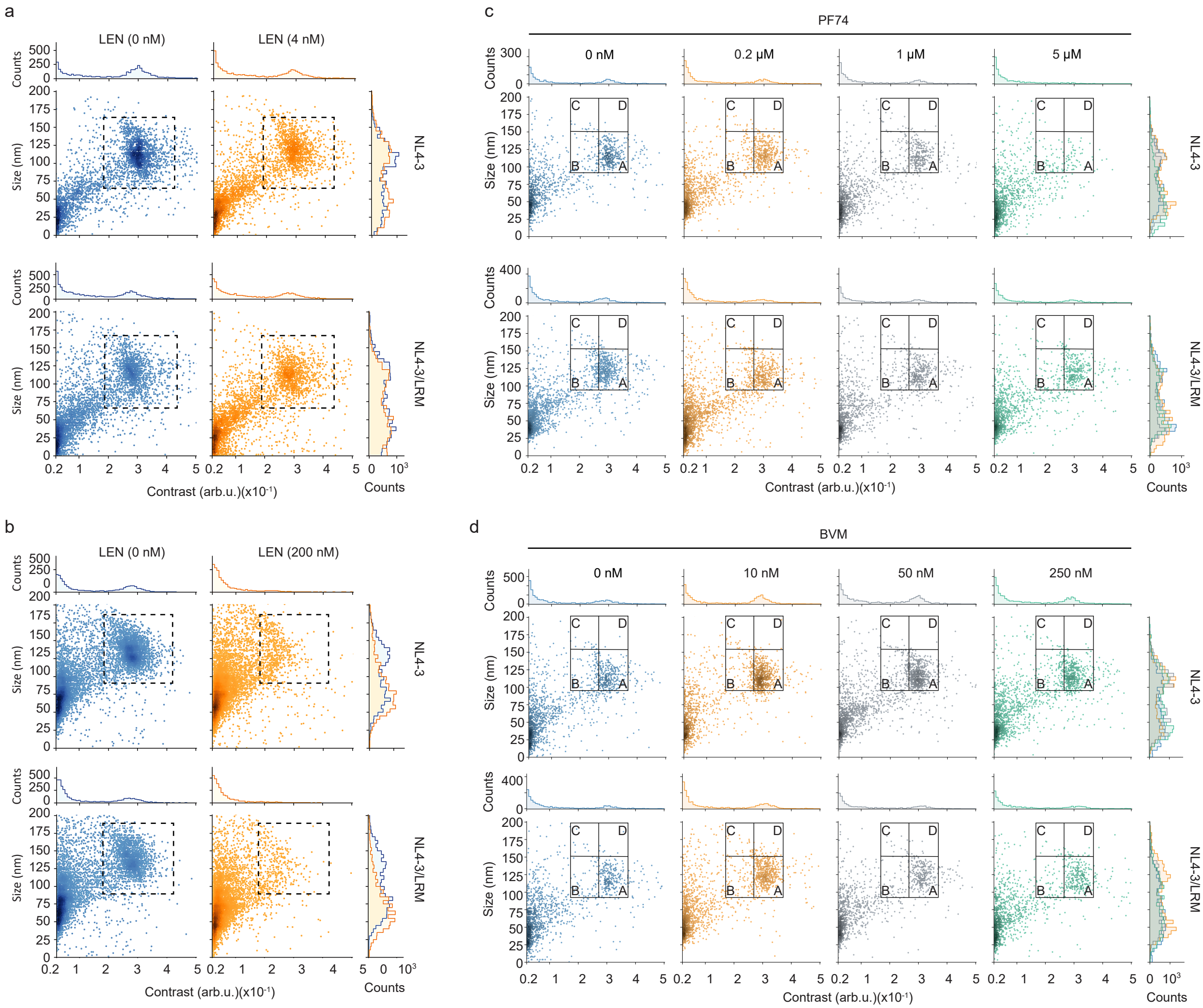

### Supplemental Figure 9

Extended data figure 9

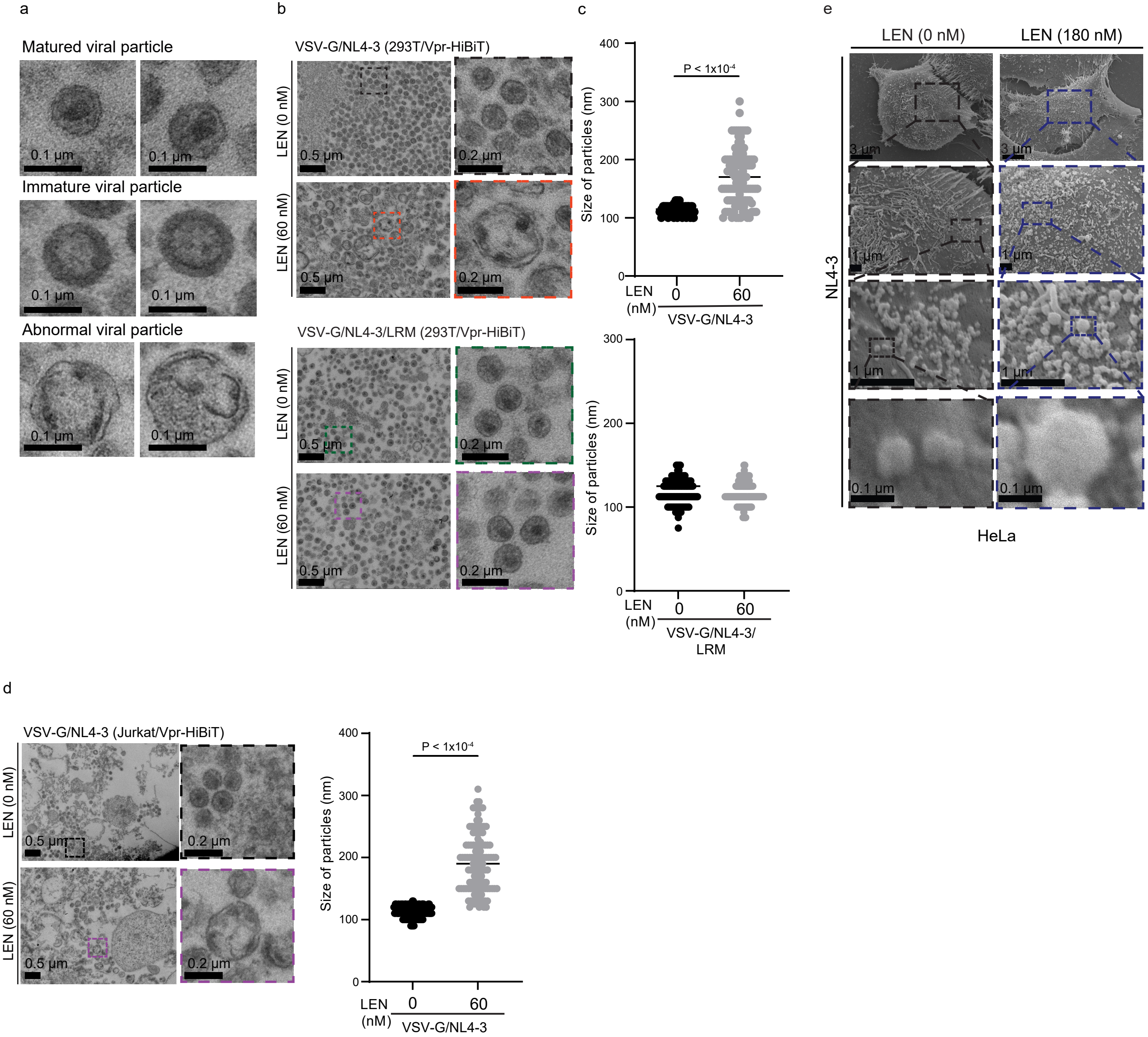

### Supplemental Figure 10

Extended data figure 10

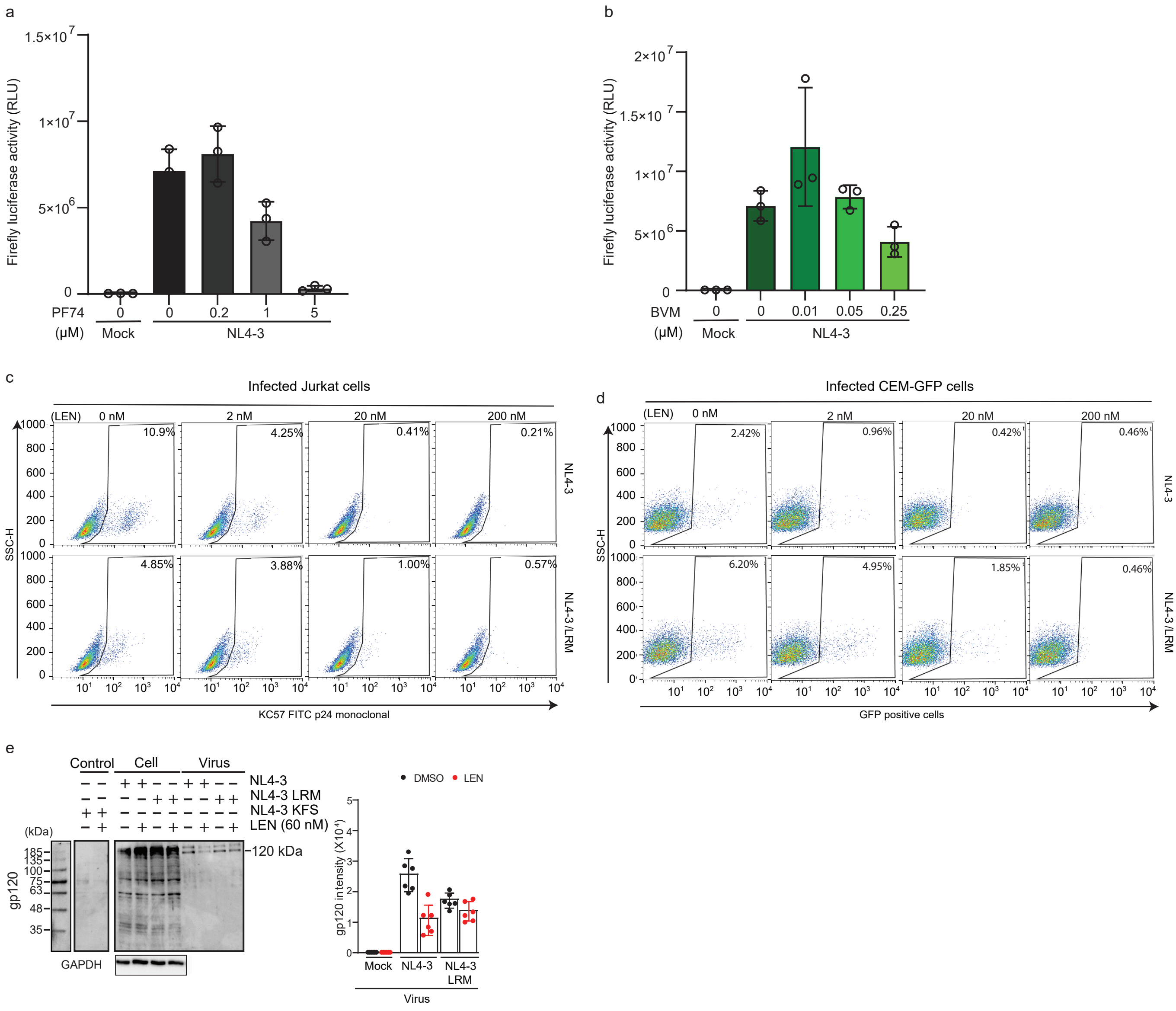
