## Supplemental Table 1 for "Lenacapavir binding to immature Gag triggers the emergence of giant HIV-1 virions"

Supplementary table 1

| Number | Clone Name | Sub-type | Fold inhibition | Rel. infectivity Inhibition (%) |
|---|---|---|---|---|
| 1 | NL4-3 | B | 1.88 | 99.96±0.03 |
| 2 | NL4-3 LRM | B | 1.53 | 15.13±3.70 |
| 3 | pWITO.C/2474 | B | 1.89 | 99.87±0.14 |
| 4 | pCH040.c/2625 | B | 1.5 | 99.25±0.82 |
| 5 | pCH058.c/2960 | B | 1.28 | 99.94±0.06 |
| 6 | pCH077.t/2627 | B | 1.43 | 99.94±0.06 |
| 7 | pCH106.c/2633 | B | 1.38 | 99.83±0.18 |
| 8 | pTHRO.c/2626 | B | 1.88 | 99.83±0.18 |
| 9 | pTRJO.c/2851 | B | 0.7 | 100 |
| 10 | pSUMA.c/2821 | B | 1.25 | 97.67±2.55 |
| 11 | 90CF402 | A/E | 1.61 | 93.41±2.55 |
| 12 | CH200C | C | 1.3 | 99.91±2.58 |
