## Supplemental Table 2 for "Lenacapavir binding to immature Gag triggers the emergence of giant HIV-1 virions"

Supplementary Table 2.

| **Viral Particle Classification** | **Characteristics** |
|---|---|
| Matured | • Single core which represents dense area observed and clearly demarcated.<br>• Two or one viral envelop surrounding the dense area.<br>• Viral structure is clearly defined.<br>• Size of particle ranges between 100-130nm. |
| Immature | • There are no dense areas centrally.<br>• Viral enveloped are clearly demarcated. |
| Abnormal | • Two or more dense areas representing the viral core are observed in one virion.<br>• Viral envelopes are not rounded as observed in mature viral particles.<br>• Viral particle structures are not clearly defined. |
